# Analysis of leaf microbiome composition of near-isogenic maize lines differing in broad-spectrum disease resistance

**DOI:** 10.1101/647446

**Authors:** Maggie R. Wagner, Posy E. Busby, Peter Balint-Kurti

**Author notes:** Correspondence and requests for materials should be addressed to M.R.W. *Current address:* Department of Ecology and Evolutionary Biology, University of Kansas, 1200 Sunnyside Avenue, Lawrence, KS, 66045, USA.

## Abstract

- Plant genotype strongly affects disease resistance, and also influences the composition of the leaf microbiome. However, these processes have not been studied and linked in the microevolutionary context of breeding for improved disease resistance. We hypothesized that broad-spectrum disease resistance alleles also affect colonization by non-pathogenic symbionts.
- Quantitative trait loci (QTL) conferring resistance to multiple fungal pathogens were introgressed into a disease-susceptible maize inbred line. Bacterial and fungal leaf microbiomes of the resulting near-isogenic lines were compared to the microbiome of the disease-susceptible parent line at two timepoints in multiple fields.
- Introgression of QTL from disease-resistant lines strongly shifted the relative abundance of diverse fungal and bacterial taxa in both 3-week-old and 7-week-old plants. Nevertheless, the effects on overall community structure and diversity were minor and varied among fields and years. Contrary to our expectations, host genotype effects were not any stronger in fields with high disease pressure than in uninfected fields, and microbiome succession over time was similar in heavily infected plants and uninfected plants.
- These results show that introgressed QTL can greatly improve broad-spectrum disease resistance while having only limited and inconsistent pleiotropic effects on the leaf microbiome in maize.

## Introduction

Phyllosphere microbiomes—the communities of bacteria and fungi living on and in plant leaves—profoundly affect the health of their plant hosts and the entire ecosystem (Lindow & Brandl, 2003; Vorholt, 2012; Laforest-Lapointe *et al*., 2017). Leaf-dwelling microbes can interfere with the exchange of gases and plant-derived volatiles (Bringel & Couée, 2015), alter patterns of herbivory (Clay, 1990; Humphrey *et al*., 2014), participate in nitrogen cycling (Murty, 1984; Papen *et al*., 2002; Fürnkranz *et al*., 2008), and influence drought resistance (Schardl *et al*., 2004; Rodriguez *et al*., 2009). Microbial symbionts are also noted for their role in disease resistance; manipulation of the phyllosphere microbiome can directly affect disease susceptibility in various species including tomato, poplar, wheat, and *Arabidopsis thaliana* (Massart *et al*., 2015; Busby *et al*., 2016; Ritpitakphong *et al*., 2016; Berg & Koskella, 2018). Despite the importance of leaf microbes to plant health, little is known about whether they are affected by systematic changes in host genotype, such as those introduced by crop breeders.

Previous studies of microbiome heritability have compared distantly related genotypes to each other, or mutated genes to the wild type (Bodenhausen *et al*., 2014; Horton *et al*., 2014; Ritpitakphong *et al*., 2016; Wagner *et al*., 2016; Wallace *et al*., 2018). Here, we take a new approach by using germplasm from a real breeding experiment to compare leaf microbiome composition before and after the introgression of quantitative trait loci (QTL) from disease-resistant maize lines into disease-susceptible lines. We designed our study to (1) test whether systematic genetic changes commonly used in breeding programs have the potential to alter crop microbiomes, and (2) to disentangle the relationships between host genotype, disease resistance, and leaf-associated microbes.

The ecological, physiological, and molecular mechanisms by which the microbiome influences disease resistance are complex and poorly understood. For instance, in *A. thaliana*, the foliar community did not directly inhibit the pathogen *Botrytis cinerea* but still conferred resistance via an unknown interaction with the plant host (Ritpitakphong *et al*., 2016). Inoculation with individual fungal endophytes substantially reduced symptoms of *Melampsora* rust infection in *Populus trichocarpa*, but other endophytes had no effect or even increased disease severity (Busby *et al*., 2016). And in tomato, the ability of the phyllosphere microbiome to improve resistance to *Pseudomonas syringae* depended on the nutrient status of the plant (Berg & Koskella, 2018). These examples illustrate the need for further investigation of the links between pathogens, the rest of the leaf microbiome, and their shared host.

One potential link between disease resistance and the microbiome is a shared sensitivity to plant genotype, which largely determines the plant phenotype. Host phenotype, in turn, determines the habitat available to both pathogenic and non-pathogenic microbes. Several studies have detected host genetic variation affecting features of the phyllosphere microbiome either among or within plant species (Sapkota *et al*., 2015; Wagner *et al*., 2016; Wallace *et al*., 2018), but most of the plant genes and traits that shape microbiome composition remain unknown. In laboratory settings, mutations in cuticle synthesis genes also affect the composition of foliar bacterial communities (Bodenhausen *et al*., 2014; Ritpitakphong *et al*., 2016), and salicylic acid signaling and glucosinolate biosynthesis genes can alter root microbiome composition (Bressan *et al*., 2009; Lebeis *et al*., 2015). A genome-wide association study of field-grown *A. thaliana* revealed that genes affecting cell wall traits, defense-response pathways, and trichome development were overrepresented among the candidate genes at quantitative trait loci (QTL) affecting foliar microbiome composition (Horton *et al*., 2014). In poplar, down-regulation of a key enzyme in the lignin biosynthetic pathway dramatically changed the composition of endophyte communities in leaves, stems, and roots (Beckers *et al*., 2016). In addition, evidence is mounting that the plant innate immune system is centrally involved in regulating microbial symbionts (Hacquard *et al*., 2017).

Some of the plant traits implicated in microbiome variation have also been implicated in quantitative disease resistance (QDR), or partial resistance to one or more pathogens (Poland *et al*., 2009; Niks *et al*., 2015; Beckers *et al*., 2016; Yang *et al*., 2017). For example, salicylic acid is a critical hormonal regulator of defense responses (Loake & Grant, 2007); and while the leaf cuticle can be a physical barrier to pathogens and a reservoir for antimicrobial compounds, it also can be recognized and used by pathogens to stimulate invasion (Martin, 1964; Bessire *et al*., 2007; Kachroo & Kachroo, 2009; Serrano *et al*., 2014). QDR is a valuable target for crop improvement for several reasons. Compared to the immunity conferred by large-effect resistance (or “R”) genes, QDR is generally more difficult for pathogens to overcome via co-evolution (St Clair, 2010). In addition, unlike the highly specific R-genes, QDR genes can be effective against several pathogens (Wisser *et al*., 2011; Wiesner-Hanks & Nelson, 2016; Yang *et al*., 2017). The resulting broad-spectrum protection, or multiple disease resistance (MDR), is desirable when several pathogens are present or disease pressures are unpredictable.

By definition, MDR loci affect colonization success of multiple pathogenic microorganisms; therefore, we hypothesized that they might also influence other microbiome members. MDR is usually a quantitative plant trait underlain by a large number of relatively small-effect genes, likely with diverse functions. Although a few MDR genes have been identified (Krattinger *et al*., 2009; Wiesner-Hanks & Nelson 2016; Sucher *et al*., 2017), most of the mechanisms underlying MDR remain unknown. Despite this, systematic breeding methods such as controlled crosses and recurrent selection enable genetic improvement of this complex trait.

We used germplasm from an MDR breeding program to test whether QTL introgressed from disease-resistant lines have pleiotropic effects on the maize leaf microbiome. We compared the foliar microbiomes of improved and unimproved maize lines in several fields, at early- and late-season timepoints, both with and without pathogen infection. Our data enabled us to test several hypotheses. First, because these MDR lines were selected for resistance to three different fungal pathogens (Lopez-Zuniga et al., 2019; Martins *et al*., 2019), we hypothesized that the introgressed alleles would have stronger effects on the fungi than bacteria. Second, because these loci have known effects on disease resistance, we hypothesized that their effects on the microbiome would be stronger in environments with higher disease pressure (Figure 1). Finally, we hypothesized that disease establishment would disrupt patterns of microbiome succession over the growing season. Our results suggest that introgression of QTL from disease-resistant lines can greatly improve broad-spectrum disease resistance with only limited, context-dependent side effects on the maize leaf microbiome.

**Figure 1.**
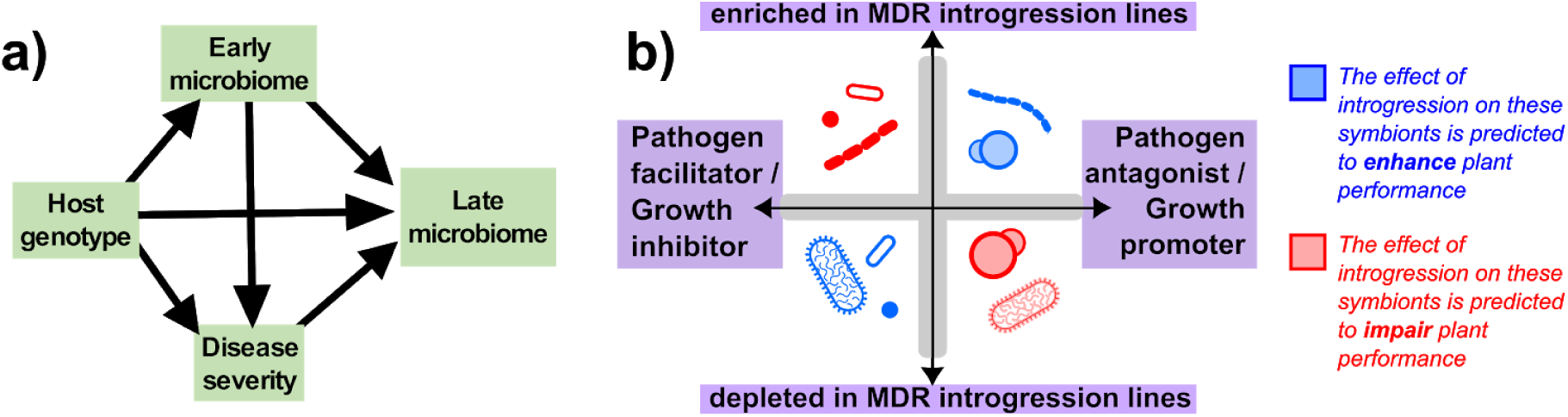
Host-pathogen-microbiome relationships involve complex interactions among all community members. **(a)** In our simplified model, host genotype can affect the late-season microbiome both directly and through cascading effects via disease resistance; for this reason, we hypothesized that MDR alleles would exert stronger effects on the microbiome when disease pressure is higher. Furthermore, host genotype could affect disease severity both directly (via immune system and other traits that impact pathogen success) and indirectly (via traits that influence early microbiome assembly, which in turn interacts with the pathogen). **(b)** The repercussions of breeding-induced changes in a microbial symbiont’s relative abundance (Fig. 4) will depend on whether it has a positive effect, negative effect, or no effect on host health. For example, if QTL introgression causes a beneficial organism to increase in abundance, or a harmful organism to decrease in abundance, the expected outcome for the host plant would be an improvement in health or performance.

## Materials and methods

### Field experimental design

To directly test whether breeding for MDR affects the foliar microbiome, we compared microbiome composition of near-isogenic plants with and without introgressed chromosome segments that conferred MDR (Lopez-Zuniga *et al*., 2019; Martins *et al*., 2019). Two MDR inbred lines (NC304 and Ki3) were crossed with H100, a highly disease susceptible line. Using single seed descent, the resulting F_1_ offspring were backcrossed three times to H100 and then self-fertilized for four generations. The resulting two populations of ∼200 BC_3_F_4:5_ near-isogenic lines (NILs) were mostly genetically identical to the recurrent elite parent (H100) but retained small chromosome segments from the donor lines. The NILs were assessed for resistance to three fungal pathogens: *Bipolaris maydis, Setosphaeria turcica*, and *Cercospora zeae-maydis*, the causative agents of the maize foliar diseases southern corn leaf blight, northern corn leaf blight, and grey leaf spot, respectively.

For this study, we selected eight NILs (four from each cross; Table S1) that had high scores for resistance to all three pathogens. The relatively strong MDR phenotypes of these NILs likely reflect larger-than-average contributions from the MDR parent genome (roughly 10% per NIL, compared to the expected 6.25% based on the breeding design; alleles from Ki3 were also more homozygous than expected (92% compared to the expected 78%) (Supporting Information Fig. S1; (Lopez-Zuniga *et al*., 2019). Within each set of four NILs there was little overlap between introgressed regions, and cumulatively the NILs carried approximately 40% of each MDR parent genome (Fig. S1). We planted these eight NILs and their parent lines in multiple fields at the Central Crops Research Station (Clayton, NC; Table S2). Replicate plots were planted in two fields in 2016, and in four fields in 2017 (Fig. 2c). Twenty kernels per line were planted per field, with the exception of the recurrent parent H100, which was planted at a replication of 30 kernels per field. Due to uneven germination, final sample sizes varied among replicates.

**Figure 2.**
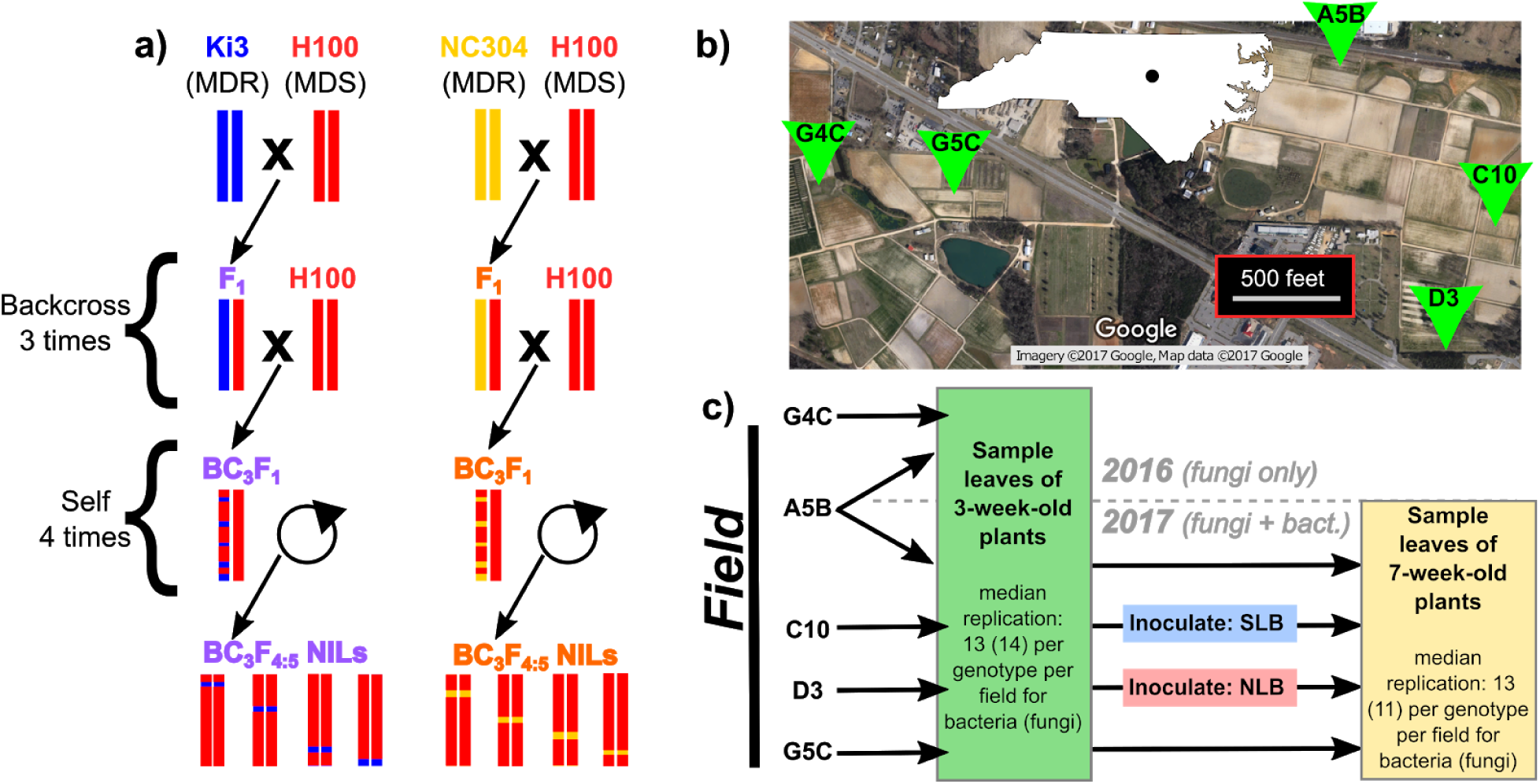
Overview of experimental design. Panel **(a)** illustrates the crossing design used to generate the eight near-isogenic lines (NILs) used in this experiment, which were mostly genetically identical to their disease-susceptible parent line H100 but which had chromosome segments introgressed from a donor line (Ki3 or NC304) that conferred multiple disease resistance (MDR). Eight NILs were planted in randomized plots along with the three parent lines. Panel **(b)** shows the locations of the replicate plots within Central Crops Research Station, Clayton, NC, USA. Map data and imagery: Google. Panel **(c)** summarizes the sampling scheme for a total of six experimental replicate plots over two years. For the pilot experiment in 2016, only a single timepoint was sampled at two fields, and only fungi were quantified. In 2017, we quantified both bacteria and fungi; plants were sampled at two timepoints in four fields, two of which were inoculated with the causal agents of either southern leaf blight (SLB; causal agent *Cochliobolus heterostrophus*) or northern leaf blight (NLB; causal agent *Setosphaeria turcica*).

To reduce microbial inoculum from kernel surfaces, we soaked kernels in 3% hydrogen peroxide for two minutes and rinsed them in distilled water immediately before planting. In each field, plants were randomly arranged in five to six adjacent rows of 40 to 50 plants each, spaced 12 inches apart. To reduce edge effects, we surrounded all experimental plots with two rows of border plants. All plots were maintained using standard agronomic conditions for rainfed maize. All fields were separated by <2 km and had similar soil types but different crop rotation histories (Fig. 2b; Table S2).

### Pathogen inoculation and disease scoring

In 2017, we explored the effects of pathogen invasion on foliar microbiomes in maize by inoculating one-month-old plants in two of the four fields. Plants in field “C10” were inoculated with *Cochliobolus heterostrophus* (the causal agent of southern leaf blight); plants in field “D3” received *Setosphaeria turcica* (the causal agent of northern leaf blight), and plants in the other two fields received no inoculation (Fig. 2c). Inoculations were performed by incubating sterilized sorghum grains in pathogen cultures, and then dropping infected grains into the whorl of each plant (Sermons & Balint-Kurti, 2018). Approximately 2 weeks after inoculation, we visually scored symptom severity of all inoculated individuals. Northern leaf blight symptoms were scored by estimating the percentage of each leaf damaged by lesions, and then averaging these scores for each plant. Southern leaf blight symptoms were scored for entire plants on a scale from 1 (complete leaf mortality) to 9 (asymptomatic) (Lopez-Zuniga *et al*., 2019). Disease scores were recorded using the Field Book application (Rife & Poland, 2014).

### Sample collection

In both 2016 and 2017, we collected leaf samples for microbiome quantification when plants were 3 weeks old. In 2017 only, we sampled leaves again when plants were 7 weeks old (i.e., 3 weeks after pathogen inoculations). The increase in experimental scope between years reflected an increase in available resources. For all sample collections, we used a standard hole punch to remove three discs evenly spaced from the base to the tip of a single leaf. For the early timepoint, we sampled the third leaf; in cases where the third leaf was too small or too damaged (<5% of plants), we sampled the second or fourth leaf instead. For the second timepoint we sampled the oldest leaf that was at least 50% green and was not touching the soil, because the microbiomes of older leaves are more likely to reflect host-driven processes than younger leaves, which are in earlier stages of microbiome assembly and more prone to stochastic influences (Maignien *et al*., 2014). We selected green tissue and avoided lesions because we were primarily interested in direct genotype effects on non-pathogenic microbial symbionts, rather than microbiome responses to differences in pathogen abundance (Fig. S2); leaves with insufficient green tissue were not sampled. Leaf discs were collected into sterile tubes and stored on ice until they could be transferred into −20°C for storage. Tools were rinsed in 70% ethanol between samples to reduce transfer of microbes among plants.

### DNA extraction, library preparation, and sequencing

To remove loosely associated microbes from leaf surfaces, we vortexed leaf discs in sterile water for 10 s on maximum speed and then shook them dry before freezing them at −80°C. Lyophilized leaf discs were randomly arranged into 96-well plates and powdered using a Retsch MM301 mixer mill (1 minute at 25 Hz). Several wells were left empty as a negative control; to several others we added a mock microbial community as a positive control (ZymoBiomics Microbial Community Standard, Zymo Research, Irvine, CA, USA). We extracted DNA using the Synergy 2.0 Plant DNA Extraction Kit (OPS Diagnostics, Lebanon, NJ, USA) following the manufacturer’s instructions, except that we doubled the length of the bead-beating step.

We generated amplicon libraries separately for bacteria and fungi using a two-PCR-step approach. First, we amplified 16S-v4 and ITS1 using the standard primer pairs 515f/806r for bacteria and ITS1f/ITS2 for fungi. Primers included upstream “frameshift” stretches of 3 to 6 random nucleotides to increase library complexity, plus a binding site for universal Illumina adaptors. Each 10-uL reaction included 0.4 uL of each primer (10 uM), 5 uL of 5Prime HotMasterMix (Quanta Bio, Beverly, MA, USA), 1.5 uL of template DNA, and 0.15 uL PNA PCR-blocker to reduce amplification of host plastid sequence (for bacterial libraries only; (Lundberg *et al*., 2013). The PCR program for fungal libraries included an initial 2-minute denaturation at 95°C; 27 cycles of 20-second denaturation at 95°C / 20-second primer annealing at 50°C / 50-second extension at 72°C; and a final 10-minute extension at 72°C. The PCR program for bacterial libraries was identical except that the primer annealing step was at 52°C and was preceded by a 5-second PNA annealing step at 78°C. The PCR products were cleaned by adding 7 uL of magnetic SPRI bead solution, washing magnet-bound DNA twice with 70% ethanol, and eluting in 10 uL ultrapure water.

The second PCR step added dual-indexed universal Illumina adaptors. The forward and reverse primers consisted of (from 5’ to 3’) the P5 or P7 adaptor sequence (respectively), a unique 8-bp index, and a binding site to enable annealing to amplicon sequences. PCR conditions were the same as above, except that only 8 cycles were performed and 1 uL of the first-step PCR product was used as the template. We then pooled 1 uL from each reaction to create separate pools for fungi and bacteria, which we purified by adding magnetic bead solution at a ratio of 0.8:1 (v/v), washing twice with 70% ethanol, and eluting DNA in ultrapure water. Aliquots of the fungal and bacterial pools were combined at equimolar concentrations.

The final combined pool derived from the 2017 samples was sequenced at 1,344-plex on an Illumina HiSeq2500 machine in Rapid Run mode (250 bp paired-end reads). To increase library complexity, a 5% phiX spike-in was added prior to sequencing. Because this first sequencing run yielded ample ITS sequence but low coverage of 16S amplicons, we sequenced the 16S amplicon pool again on the HiSeq platform and on the MiSeq using V2 chemistry (250 bp paired-end reads) along with the smaller pool of ITS amplicons from the 2016 samples. All sequencing was performed by the North Carolina State University Genomic Sciences Laboratory (Raleigh, NC, USA).

### Sequence processing and quality filtering

After trimming primers from raw, demultiplexed FASTQ files using _CUTADAPT_ v1.12 (Martin, 2011), we processed sequences using _DADA_2 v1.10.1 (Callahan *et al*., 2016). We required the forward and reverse 16S reads to have a maximum of 2 expected errors and no ambiguous bases, then truncated them at 220bp and 160 bp, respectively. We required the forward and reverse ITS reads to have a maximum of 1 and 2 expected errors (respectively) and no ambiguous bases but did not truncate reads to a fixed length. Error rates were inferred from 3×10^6^ reads; this was done separately for the ITS data and 16S data, and separately for each independent sequencing run. Quality-filtered reads were then de-replicated, de-noised, and merged to generate tables of amplicon sequence variants (ASVs). At this point we merged the bacterial ASV tables from the three 16S sequencing runs with each other, and also merged the fungal ASV tables from 2016 and 2017, which had been sequenced separately. After removing chimeric ASVs, we assigned taxonomy using the RDP Classifier (Wang *et al*., 2007) trained on the RDP (v.16) training set for 16S sequences and the UNITE database for ITS sequences (Cole *et al*., 2014)(Kõljalg *et al*., 2005).

We discarded ASVs without taxonomic assignment at the kingdom level and ASVs that were assigned to chloroplasts or mitochondria (“non-usable reads”). We used the mock community positive controls to determine a within-sample relative abundance threshold that removed most contaminant ASVs while retaining as much of the data as possible. This threshold (0.091% for bacteria, 0.221% for fungi) was then applied to all non-control samples. We then removed “non-reproducible” ASVs that were not observed at least 25 times in at least 5 samples (Lundberg *et al*., 2012). Together, these filtering steps reduced the final dataset to 1,502 bacterial ASVs while retaining 93.3% of the data. For fungi, the final dataset retained 548 ASVs and 90.5% of the original sequences. Finally, we excluded samples with <500 usable reads. Out of the original 1,728 fungal samples, 194 were excluded from analysis; for bacteria, 174 out of 1,315 were excluded. The number of reads remaining after all filtering steps was saved as the “sampling effort” for each sample, normalized and centered for use as a covariate in downstream analyses.

### Data analysis

We used R version 3.6.0 for all analysis, especially the packages phyloseq, tidyr, lme4, DESeq2, vegan, and lmerTest (McMurdie & Holmes, 2013; Love *et al*., 2014; Bates *et al*., 2015; Kuznetsova *et al*., 2017). When applicable, we used the false discovery rate (FDR; (Benjamini & Hochberg, 1995) to adjust *P*-values from multiple comparisons. All analyses were performed in parallel for fungi and bacteria. Original R code and raw data are available in a Zenodo repository (Wagner *et al*. 2019); raw reads are available in the NCBI Sequence Read Archive under BioProject #PRJNA565009.

We estimated alpha diversity for each sample using the Shannon and abundance-based coverage estimator (ACE) metrics, which describe community evenness and richness, respectively (Hughes *et al*., 2001). For analyses conducted at higher taxonomic levels, we consolidated ASVs into their respective genera, families, orders, classes, or phyla using the function “Phyloseq::tax_glom” (McMurdie & Holmes, 2013). For analyses that required normalization (*e*.*g*., ordination) we applied the variance-stabilizing transformation from the “DESeq2” package (Love *et al*., 2014; McMurdie & Holmes, 2014). Bray-Curtis dissimilarity of transformed count data was used to quantify community composition. When modeling the relative abundances of individual taxa using the DESeq2 package, we tested only taxa with abundances that were at least 10% of the mean taxon abundance (Wagner *et al*., 2016). For example, the mean bacterial ASV was observed 50,689 times across the full dataset; therefore, we excluded all bacterial ASVs that were observed fewer than 5,069 times from the relevant analysis. This greatly reduced the number of tests to be performed but retained most of the data; for example, across the full dataset it reduced the number of bacterial ASVs from 1502 to 576 while retaining 98.9% of all observations. We explored overall patterns of microbiome variation by performing multivariate ANOVA on the Bray-Curtis dissimilarity matrix of the full variance-stabilized dataset using the function “vegan::adonis” (Oksanen *et al*., 2018). This model included the predictor variables “Genotype”, “Rep” (i.e., field and year), “Genotype*Rep”, “Timepoint”, and “Genotype*Timepoint”.

#### Characterization of beta diversity and changes in beta diversity

In addition to overall microbiome composition, we were interested in whether QTL introgression from disease-resistant lines affected microbiome *variability*. One host genotype might be hospitable to only a small subset of microbes, whereas another may be open to colonization by a wider range of symbionts. The former host would be expected to exhibit low variability among biological replicates, whereas the latter host has the potential to exhibit higher variability among biological replicates due to stochastic and microenvironmental effects. Such a scenario would manifest as a host genotype effect on beta dispersion.

Beta dispersion for groups of samples was calculated using the function “vegan::betadisper” (Oksanen *et al*., 2018). For such analyses, samples were grouped in several different ways depending on the question being asked. For example, to ask whether beta diversity differed between early and late timepoints, we used the function “vegan::betadisper” to find a centroid location in ordination space for each timepoint, and then calculated each individual sample’s distance to its corresponding centroid. This “Distance_to_Centroid” metric could then be used to compare beta diversity of the two groups using standard statistical approaches as detailed below. We used the same approach to assess differences in beta diversity among host genotypes and between timepoints in specific fields.

These analyses of beta diversity tested whether leaf microbiomes of one group of individual plants were more homogeneous than those of another group; however, they were not meant to compare overall microbiome composition between the groups. Rather, our conclusions about differences in microbiome composition between genotypes were drawn from the multivariate ANOVA and negative binomial models described above and below.

#### Testing effects of MDR alleles on the juvenile and adult maize microbiomes

Next, we tested the hypothesis that the introgression of MDR alleles altered microbiome composition. We conducted these analyses separately for young plants measured three weeks after planting (the “early” timepoint) and seven weeks after planting (the “late” timepoint). For each timepoint we performed multivariate ANOVA of Bray-Curtis dissimilarity, using a model that included “Genotype”, “Rep”, and their interaction as predictor variables. Because we were specifically interested in contrasting MDR genotypes to the susceptible line H100 (Fig. 2a), we repeated this analysis ten times; each time we subset the data to include only H100 and one MDR genotype. *P-*values from this analysis were corrected for multiple comparisons using the false discovery rate; FDR < 0.05 was considered statistically significant (Benjamini & Hochberg, 1995).

We took a similar approach to test whether QTL introgression from disease-resistant lines altered alpha diversity and beta diversity. We modeled ACE, Shannon, and beta diversity (*i*.*e*., distance to the centroid for the corresponding Genotype within each Rep) using separate linear mixed-effects models with “Genotype”, “Rep”, and their interaction as fixed-effect predictors. ACE diversity was natural log-transformed to improve homoscedasticity. Standardized sequencing depth and a “Plate” random-intercept term were also included as nuisance variables to control for variation in sampling effort among samples and batch effects during DNA extraction and library preparation. Post-hoc Dunnett *t-*tests (Dunnett, 1955) were used to directly contrast each MDR genotype to H100 within each Rep while controlling the family-wise error rate. Finally, to determine which microbial taxa responded to host genotype, we fit negative binomial models to counts of individual ASVs, genera, families, orders, classes, and phyla, using “Genotype”, “Rep”, and their interaction as predictor variables. For these analyses, H100 was set as the reference genotype, so that the coefficients from the model described contrasts between MDR lines and the disease-susceptible control. *P-*values were adjusted to correct for multiple comparisons (across all taxa tested, in ten MDR genotypes, in multiple experimental replicates) at FDR < 0.05.

#### Investigating the effect of disease severity on seasonal microbiome dynamics

We analyzed the effect of pathogen invasion on the microbiome by comparing microbiome succession between timepoints, (1) in inoculated versus uninoculated fields, and (2) as a function of infection severity at the individual plant level within each field. First, we performed a partial constrained distance-based redundancy analysis (based on the Bray-Curtis dissimilarity metric) to characterize the overall community response to Timepoint*Field interactions after controlling for sequencing depth. To assess statistical significance of this interaction, we used permutation tests to compare this model to an alternative model containing only the Timepoint and Field main effects. To determine which taxa drove this interaction, we used the DESeq2 package to fit negative binomial models for counts of individual ASVs, genera, families, orders, classes, and phyla in response to the Timepoint*Field interaction; likelihood ratio tests were used to compare these to alternative models with only the Timepoint and Field main effects. To investigate how disease establishment affected alpha and beta diversity at the field level from early season to late season, we calculated each individual plant’s change in Shannon diversity and in Distance_to_Centroid between timepoints (centroid calculated for each Field at each Timepoint). We then fit linear mixed models to these calculated values with “Field” as a fixed-effect predictor. Standardized sequencing depth and a “Plate” random-intercept term were also included as nuisance variables to control for variation in sampling effort and batch effects. Statistical significance was assessed using ANOVA with Type III sums of squares and Satterthwaite’s approximation for denominator degrees of freedom. We used Tukey’s Honest Significant Difference test to compare the early-to-late changes in alpha and beta diversity among fields while controlling the family-wise error rate. Second, within each inoculated field, we regressed each plant’s change in alpha diversity and in community composition (i.e., Bray-Curtis dissimilarity between the two timepoints) against symptom severity.

## Results

The final fungal dataset included 546 ASVs and 1,533 samples from 6 replicate plots over two years (2016-2017). The bacterial dataset included 1,502 ASVs and 1,141 samples from 4 replicate plots in 2017 only. The 2017 data included two timepoints: early (3-week-old plants) and late (7-week-old plants), whereas the 2016 data represented the early timepoint only (Fig. 2c). Median replication ranged from *N*=11 to *N*=14 per genotype per replicate (Fig. 2c). The median sequencing depth per sample was 29,283 for fungi and 97,955 for bacteria.

Bacterial microbiomes were structured largely by timepoint, which explained 17.6% of the variation in community composition (Fig. 3; Table 1). Experimental replicate (i.e., field) and host genotype each explained only about 3% of the variation. At the early timepoint, communities were dominated by *Pantoea* spp. (53.5% relative abundance) followed by *Herbaspirillum* spp. (12.3%). However, four weeks later, the relative abundances of these groups had declined sharply to 4.4% and 2.0%, respectively. The dominant bacterial members of the adult maize leaf microbiome belonged to the genera *Sphingomonas* (38.9%) and *Methylobacterium* (29.2%; Table S3).

**TABLE 1.**
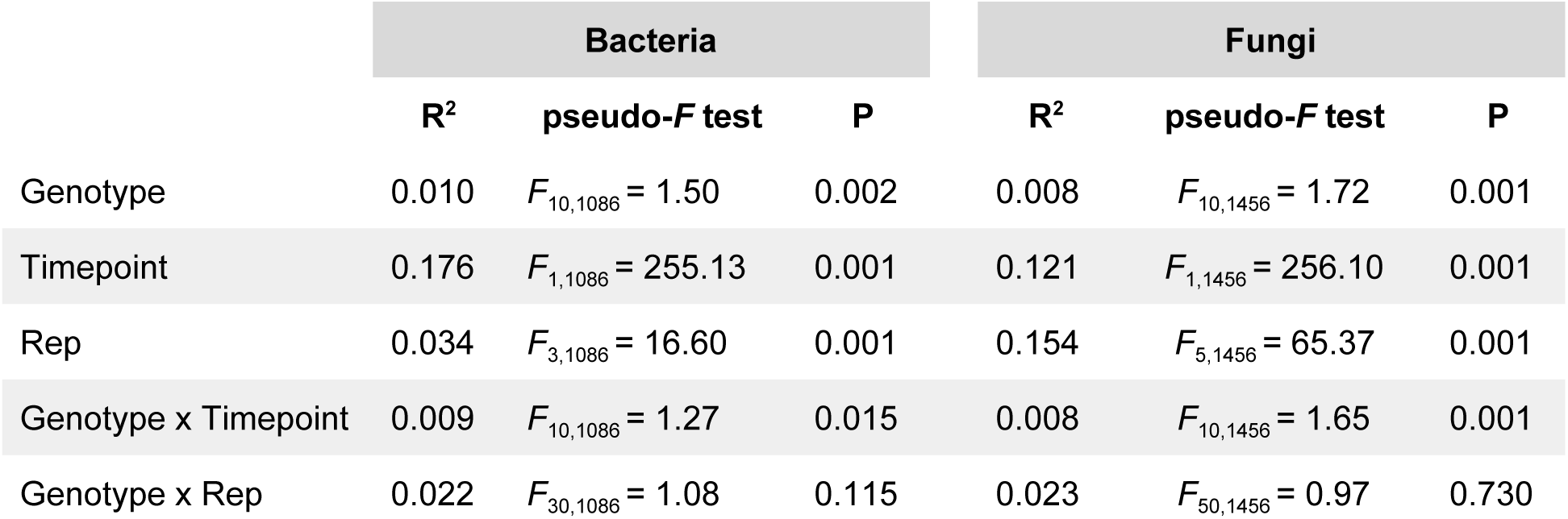
Results of permutational MANOVA for fungal and bacterial community composition in the leaves of maize plants. *P*-values are based on 999 permutations of the Bray-Curtis dissimilarity matrix calculated from variance-stabilized amplicon sequence variant (ASV) tables.

**Figure 3.**
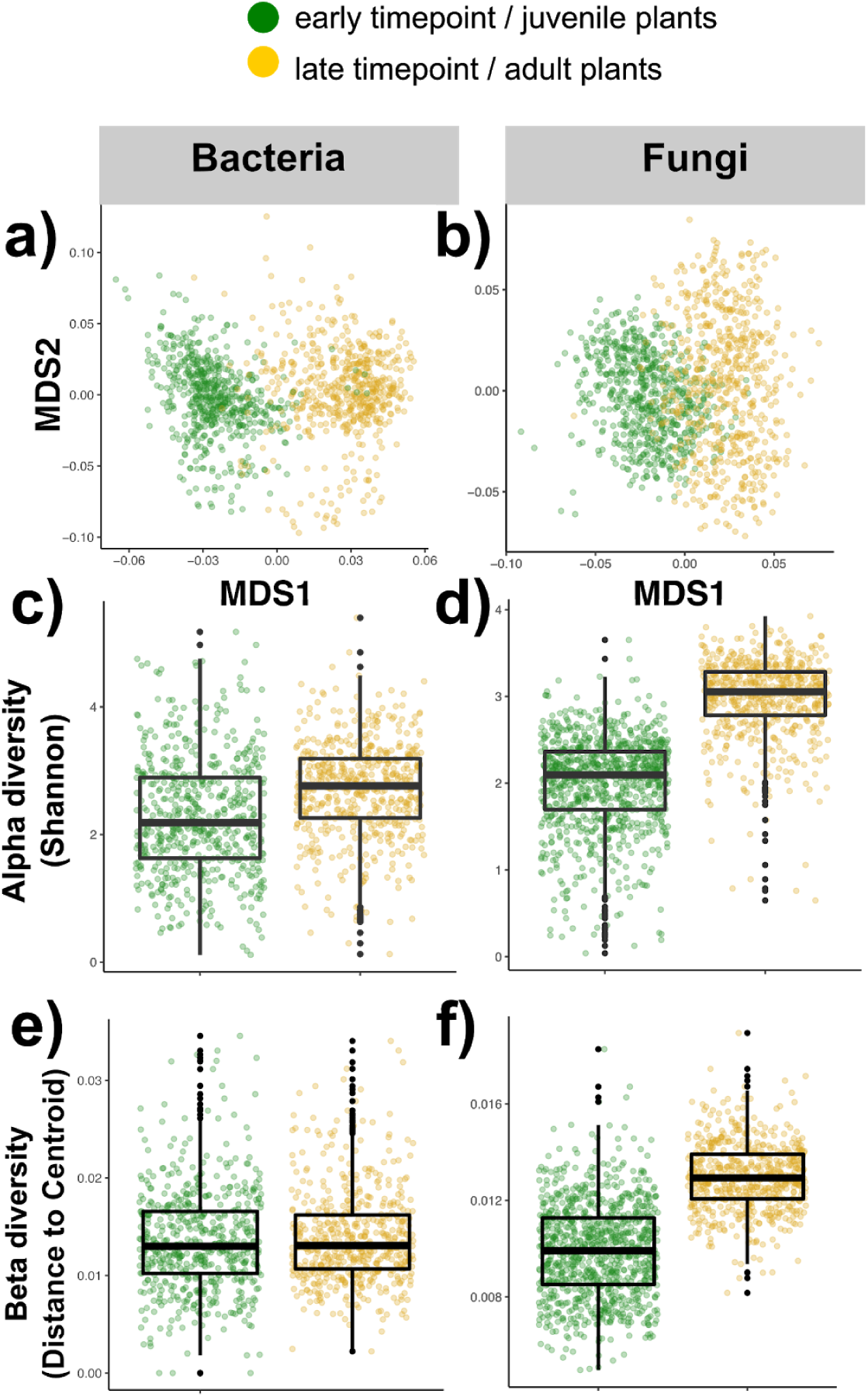
Maize leaf microbiomes shifted dramatically between 3 weeks and 7 weeks after planting. **(a-b)** Overall microbiome composition shifted strongly between timepoints. MDS1 and MDS2 are the two major axes of variation after ordination of the Bray-Curtis dissimilarity matrix using non-metric multidimensional scaling, i.e., numerical summaries of community composition. Each point represents one leaf sample; points separated by smaller distances in MDS space indicate samples with more similar microbiomes. **(c-d)** On average, alpha diversity was higher at the late timepoint than the early timepoint. The top, middle, and bottom lines of the boxes mark the 75th percentile, median, and 25th percentile, respectively; box whiskers extend 1.5 times the interquartile range above and below the box. **(e-f)** Beta diversity (*i*.*e*., variation among samples) was stable over time for bacteria, but increased for fungi between timepoints. Boxplot statistics are the same as in panels (c-d).

In contrast, fungal communities were strongly shaped by experimental replicate (i.e., field and year; Table 1); however, timepoint became the dominant predictor when data from 2016 were excluded, indicating that differences between years contributed to this result (Fig. S3). In 2016 the most abundant fungal genus in seedling leaves was *Sporobolomyces* (31.7% relative abundance) followed by *Epicoccum* (12.7%). The following year, the same genera were again the two most common in young leaves, although in the opposite order (*Epicoccum* 24.7%, *Sporobolomyces* 8.3%). In older plants, *Epicoccum* remained the most abundant genus, despite declining to 9.8% relative abundance. Overall, there was a high degree of overlap in the most abundant genera within leaf microbiomes of seedlings in 2016 and 2017 (Table S3). However, one third of ASVs--representing a diverse range of genera--changed significantly in relative abundance between years (Fig. S4).

We detected host genetic effects on overall composition of both bacterial and fungal microbiomes, as well as an interaction between host genotype and timepoint (Table 1); we explore these results in more detail below. On average, alpha diversity of both kingdoms was higher in seven-week-old plants relative to three-week-old plants (Fig. 3c-d). In contrast, beta diversity (*i*.*e*., variation among samples) of bacterial communities did not change between timepoints, whereas beta diversity of fungal communities increased (Fig. 3e-f).

### In juvenile plants, QTL introgression altered the relative abundances of diverse taxa but not overall community structure

First, we investigated whether the introgression of QTL from MDR genotypes altered microbiome composition in the leaves of young maize plants, before the establishment of disease. For these analyses we used data from both years, but included only the data from the early timepoint (3 weeks after planting). Alpha diversity of both bacterial and fungal leaf microbiomes, measured using the ACE metric, varied among genotypes (ANOVA, Genotype x Rep, *P* = 0.070 and *P* = 0.0038 respectively; Table S4). However, the strength and direction of this effect varied across experimental replicates. In some replicates, the NILs deviated from H100 in the same direction as the MDR parent lines, consistent with the hypothesis that the introgressed MDR alleles affect both disease resistance and early microbiome diversity. In others, however, there was no apparent genetic variation at all (Fig. S5). This result suggests that host genotype interacts with the environment in complex ways to influence the taxonomic diversity of leaf-associated microbial communities in maize. Tests of beta diversity--*i*.*e*., variation in microbiome composition among individuals of the same genotype--showed similar patterns. Beta diversity of both fungal and bacterial communities varied among genotypes, but the direction and strength of the effect were inconsistent among experimental replicates (Table S5; Fig. S6).

In addition to alpha and beta diversity, we investigated the effects of QTL introgression from MDR lines on overall community composition using permutational MANOVA. Only one of the two disease-resistant parent lines (and none of the NILs) differed from H100 in bacterial microbiome composition (Table 2). Similarly, fungal community composition did not differ between any of the MDR lines and H100, contradicting our hypothesis that these loci would have stronger and more consistent effects on fungi than bacteria. Nevertheless, we detected a diverse range of individual taxa that changed in relative abundance in response to MDR allele introgression. For instance, 33 fungal genera were either enriched or depleted in at least one NIL relative to the common disease-susceptible parent line H100, with effect sizes ranging from approximately 4-fold to over 1000-fold (Wald test, FDR < 0.05; Fig. 4a; Table S6). These differing patterns detected by permutational MANOVA and by negative binomial models are not necessarily contradictory; the former method can detect simultaneous shifts in a large number of species, even if most or all of those shifts are too subtle to be detected using univariate models (Anderson, 2001). Similarly, strong responses by a relatively small number of taxa--such as those observed in MDR NILs (Fig. 4)--may be missed by permutational MANOVA if the rest of the community stays relatively stable.

**TABLE 2.** Results of permutational MANOVA of fungal and bacterial community composition in maize leaves at two timepoints. Each MDR line was individually compared to the common disease-susceptible genetic background, H100. ***R***^**2**^ values are shown for the Genotype and Genotype*Rep terms of each model. For the early timepoint, the Replicate factor included variation among fields and between years; for the late timepoint, it only included variation among fields. Statistical significance was based on comparison of pseudo-*F* values after 999 permutations of the Bray-Curtis dissimilarity matrix calculated from variance-stabilized ASV tables.

| MDR line<br>(vs. H100) | Bacteria |  |  |  | Fungi |  |  |  |
| --- | --- | --- | --- | --- | --- | --- | --- | --- |
|  | Early<br>timepoint |  | Late<br>timepoint |  | Early<br>timepoint |  | Late<br>timepoint |  |
|  | Geno. | Geno.<br>x Rep | Geno. | Geno.<br>x Rep | Geno. | Geno.<br>x Rep | Geno. | Geno. x<br>Rep |
| Ki3 | **0.014\*** | 0.024 | **0.016\*** | 0.026 | 0.003 | 0.016 | 0.009<sup>‡</sup> | 0.023<sup>‡</sup> |
| DRIL32.063 | 0.009 | 0.026 | 0.012 | 0.030 | 0.005 | 0.023 | **0.011 \*** | 0.018 |
| DRIL32.095 | 0.011<sup>‡</sup> | 0.022 | 0.007 | 0.022 | 0.004 | 0.017 | 0.005 | 0.019 |
| DRIL32.134 | 0.009 | 0.021 | 0.007 | 0.023 | 0.004 | 0.019 | **0.012 \*** | 0.020 |
| DRIL32.140 | 0.007 | 0.018 | 0.008 | 0.019 | 0.002 | 0.017 | 0.007 | 0.015 |
| DRIL62.030 | 0.009 | 0.025 | 0.011 | 0.032 | 0.006 | 0.023 | **0.019 \*\*** | 0.022 |
| DRIL62.032 | 0.008 | 0.024 | 0.010 | 0.030 | 0.004 | 0.019 | 0.009<sup>‡</sup> | 0.025<sup>‡</sup> |
| DRIL62.054 | 0.009 | 0.023 | 0.008 | 0.026 | 0.005 | 0.018 | 0.008 | 0.022 |
| DRIL62.127 | 0.008 | 0.021 | 0.006 | 0.025 | 0.002 | 0.015 | 0.008 | 0.019 |
| NC304 | 0.009 | 0.028 | **0.022\*** | 0.031 | 0.008 | 0.024 | **0.021 \*\*** | **0.029\*** |

<sup>‡</sup>FDR<0.1, \*FDR<0.05, \*\*FDR<0.01

**Figure 4.**
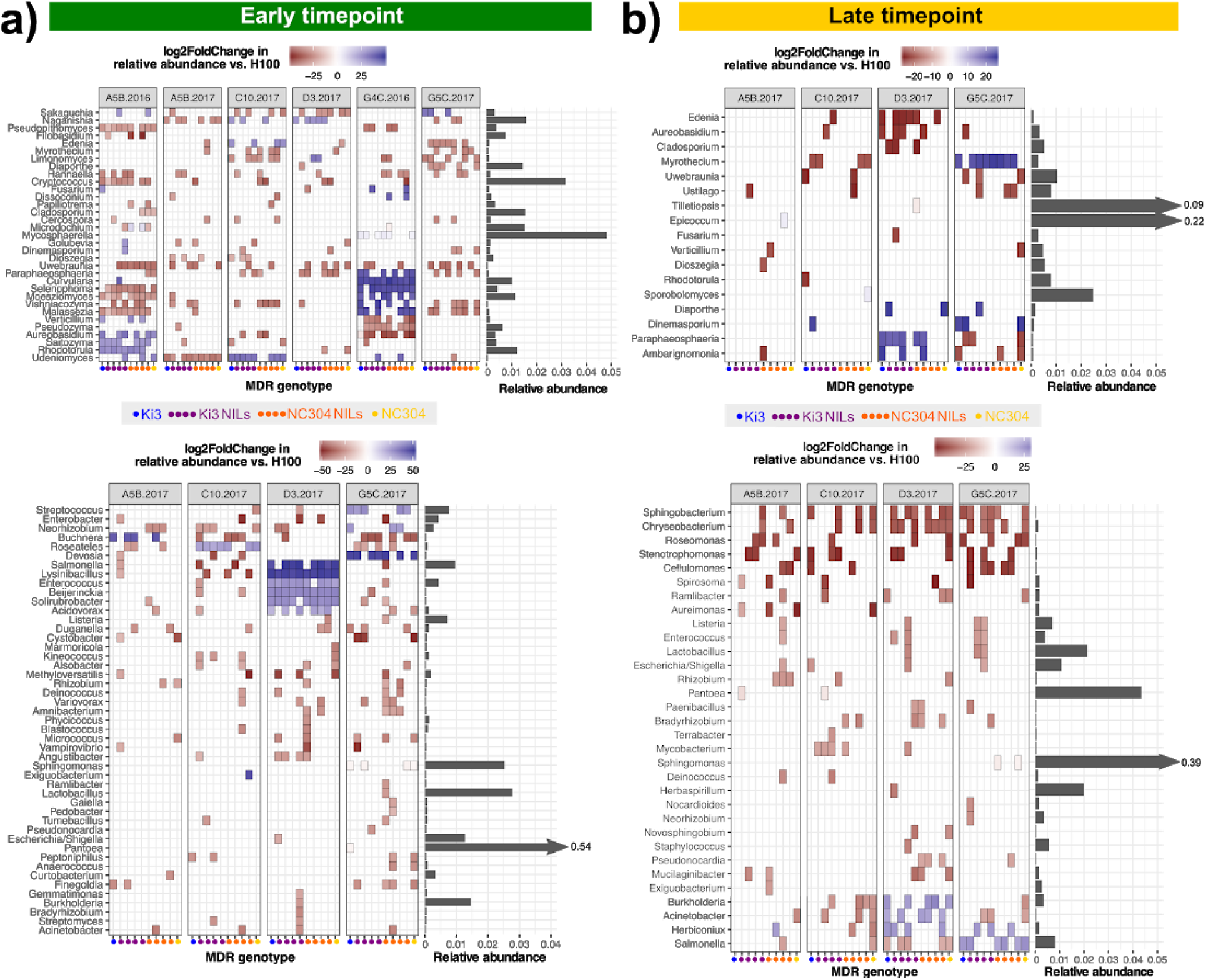
Introgression of QTL from MDR lines altered the relative abundance of diverse taxa in leaves of 3-week-old and 7-week-old maize. The enrichment/depletion of fungal (top) and bacterial (bottom) genera caused by introgression of MDR alleles into the H100 genetic background is shown for **(a)** the early timepoint/juvenile plants, and **(b)** the later timepoint/adult plants. MDR genotypes “Ki3” and “NC304” are the parent lines; the others are NILs derived from crosses between those lines and the disease-susceptible line H100 (Fig. 2). Taxa with significant decreases or increases in relative abundance (Wald test, FDR < 0.05) are shown in red or blue, respectively. The relative abundance of each genus is shown to the right of each heatplot; to improve figure clarity for the less abundant taxa, the x-axes were truncated and the relative abundances of the more common taxa are shown numerically. Fungal taxa unidentified at the genus level were excluded for clarity. Additional data on enrichment/depletion of organisms from other taxonomic levels are provided in Table S6.

Many taxa responded similarly to several introgressions. For example, several groups (including *Neorhizobium, Cryptococcus*, and *Uwebraunia*) were consistently enriched or depleted in at least five NILs. This strengthens the evidence that broad-spectrum disease resistance shares a genetic basis with the abundance of certain microbiome members, because multiple non-overlapping introgressions had similar effects on these taxa. However, the inconsistency of the microbiome response across fields and years suggests that these QTL have lower penetrance for microbiome composition than for disease resistance (Lopez-Zuniga *et al*., 2019). For instance, many other taxa (such as *Buchnera, Selenophoma, Moesziomyces, Udeniomyces*, and *Naganishia*) were strongly and consistently enriched across five or more NILs in one environment, but were consistently depleted or unaffected in other environments (Fig. 4; Table S6).

### In adult plants, introgressed QTL improved disease resistance with minimal effects on microbiome diversity

Next, we investigated whether QTL introgressed from MDR lines affected the maize leaf microbiome later in the season, seven weeks after planting. Three weeks prior to this late-season sampling, plants in two of the four fields received pathogen inoculations so that at the 7-week timepoint plants in field C10 were infected with southern leaf blight and those in field D3 were infected with northern leaf blight. We scored disease symptoms two weeks after inoculation and confirmed that resistance to both diseases was improved in all eight MDR NILs relative to the susceptible parent line H100 (Fig. 5; all *P* < 4.7e^-7^, all R^2^ > 0.70). However, we collected microbiome data only from green tissue, avoiding lesions of infected plants (Fig. S2). ASVs corresponding to the introduced pathogens (*Bipolaris maydis* and *Setosphaeria turcica*) were removed from the dataset before analysis because we were primarily interested in direct effects of MDR alleles on the non-pathogenic microbiome, rather than cascading effects on the microbiome driven by improved disease resistance.

**Figure 5.**
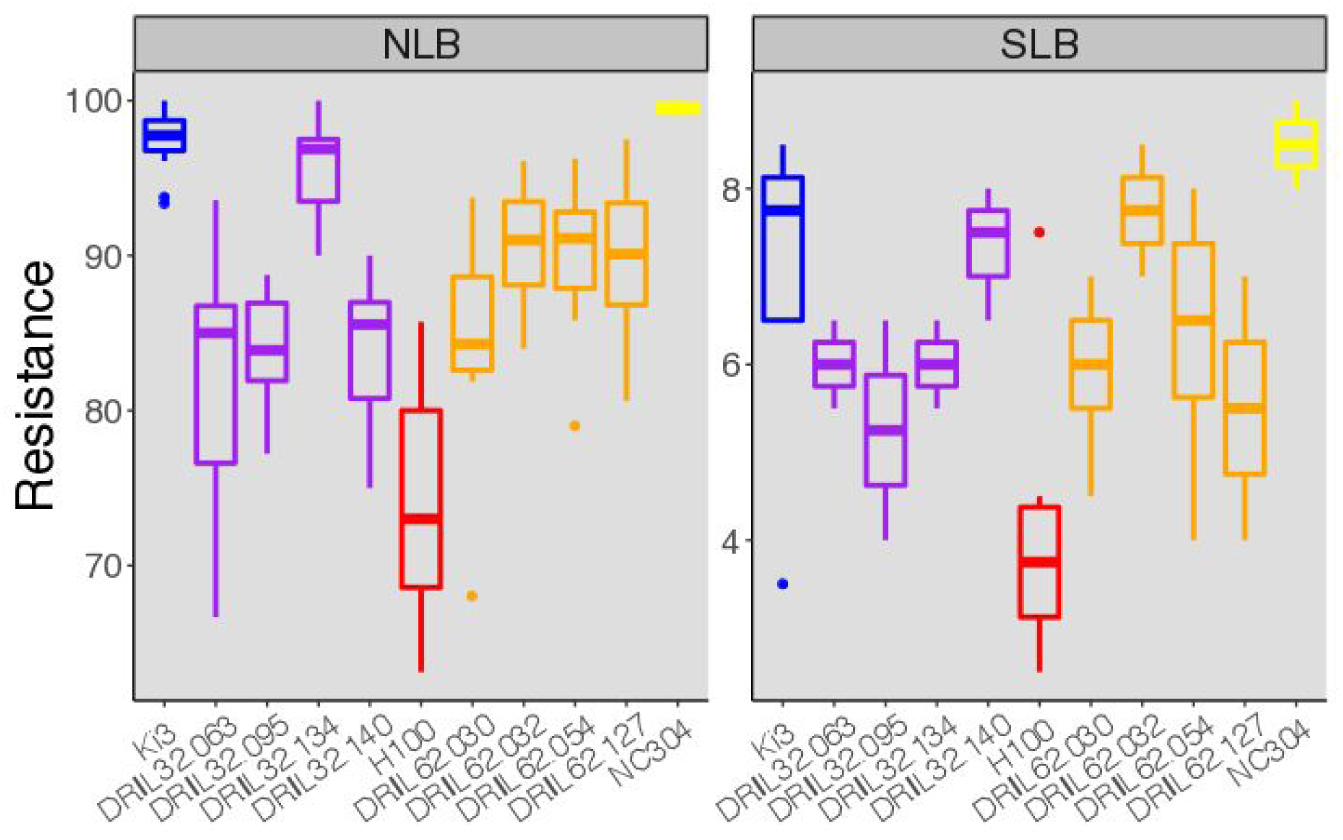
Introgression of QTL from two MDR parent lines improved resistance to northern leaf blight (NLB; left) and southern leaf blight (SLB; right) in six-week-old plants. Symptoms were scored two weeks after pathogen inoculation. The top, middle, and bottom lines of the boxes mark the 75th percentile, median, and 25th percentile, respectively; box whiskers extend 1.5 times the interquartile range above and below the box. For NLB, all comparisons to the susceptible genetic background H100 were significant at *P* < 9.3e^-4^ (*N* = 141; genotype R^2^=0.71); for SLB, all comparisons were significant at *P* < 1.6e^-7^ (*N* = 147; genotype R^2^=0.76).

Our results provided mixed support for our hypothesis that host genotype effects would be stronger at the late timepoint (after disease establishment) than the early timepoint. Permutational MANOVA showed that the introgressed QTL had stronger effects on overall community structure in the late timepoint, particularly for fungi (Table 2). For both kingdoms, genetic differences in alpha diversity were minor and were comparable between timepoints (Fig. S5). QTL introgression tended to decrease beta diversity of fungal communities only at the later timepoint, and only in the two fields that had been inoculated with pathogens. This suggests that at least some of their effects on the microbiome were mediated through their effects on disease resistance (Fig. S6). However, we detected considerably more host-genotype-sensitive taxa at the earlier timepoint (Fig. 4; Table S6). Although there was some overlap between the sets of taxa responding to introgression at the early and late timepoints, the patterns of depletion and enrichment often differed. For instance, several bacterial genera that were frequently and strongly depleted in MDR lines at the late timepoint (*Sphingobacterium, Chryseobacterium, Roseomonas, Stenotrophomonas*, and *Cellulomonas*) were not affected by the introgressions at the early timepoint (Fig. 4). This indicates that introgression-induced microbiome differences in seedlings did not generally persist throughout the growing season.

Altogether, our results indicate that QTL introgression from disease-resistant lines shifted the relative abundance of diverse bacterial and fungal taxa in the leaves of 3-week-old and 7-week-old maize plants (Fig. 4). However, the effects of these introgressions on the microbiome were much more variable among environments than their effects on disease resistance (Lopez-Zuniga *et al*., 2019; Martins *et al*., 2019). This suggests that changes in the relative abundance of potentially protective microbes is unlikely to be a major mechanism by which these particular MDR alleles confer improved disease resistance.

### Seasonal microbiome dynamics were largely insensitive to disease status

Finally, we shifted our focus away from host genotype to investigate the relationship between disease and the microbiome more closely. We hypothesized that heavy pathogen infection and disease establishment would disrupt the normal succession of maize leaf microbiomes both (1) at the whole-field level and (2) at the individual plant level. To test these hypotheses, we compared patterns of microbiome change over time in two pathogen-infected fields versus two uninfected control fields, and in heavily-infected individual plants versus less-infected individuals of the same genotype within a field.

Microbiome composition and diversity of all fields changed dramatically between three weeks and seven weeks after planting, regardless of infection status (Fig. 6). Community composition diverged among fields over time (Fig. 6a; distance-based redundancy analysis, Timepoint*Field *P* = 0.001), although this pattern was much more pronounced for fungi than bacteria. Notably, fungal communities in pathogen-inoculated fields diverged in overall composition from those in non-inoculated fields (Fig. 6a). However, the relative abundances of individual taxa generally changed in the same direction over time in all four fields (Fig. 6b). Furthermore, the average shift in relative abundance between timepoints was similar in magnitude between infected and uninfected fields (Fig. 6b-c). Similarly, temporal changes in alpha and beta diversity varied in magnitude among fields for both bacteria and fungi (Fig. 6d; ANOVA *P* < 0.05 for all); however, these differences did not correspond to disease treatment. Together, these results did not support our prediction that microbiome composition would shift more dramatically over time in environments with higher disease pressure.

**Figure 6.**
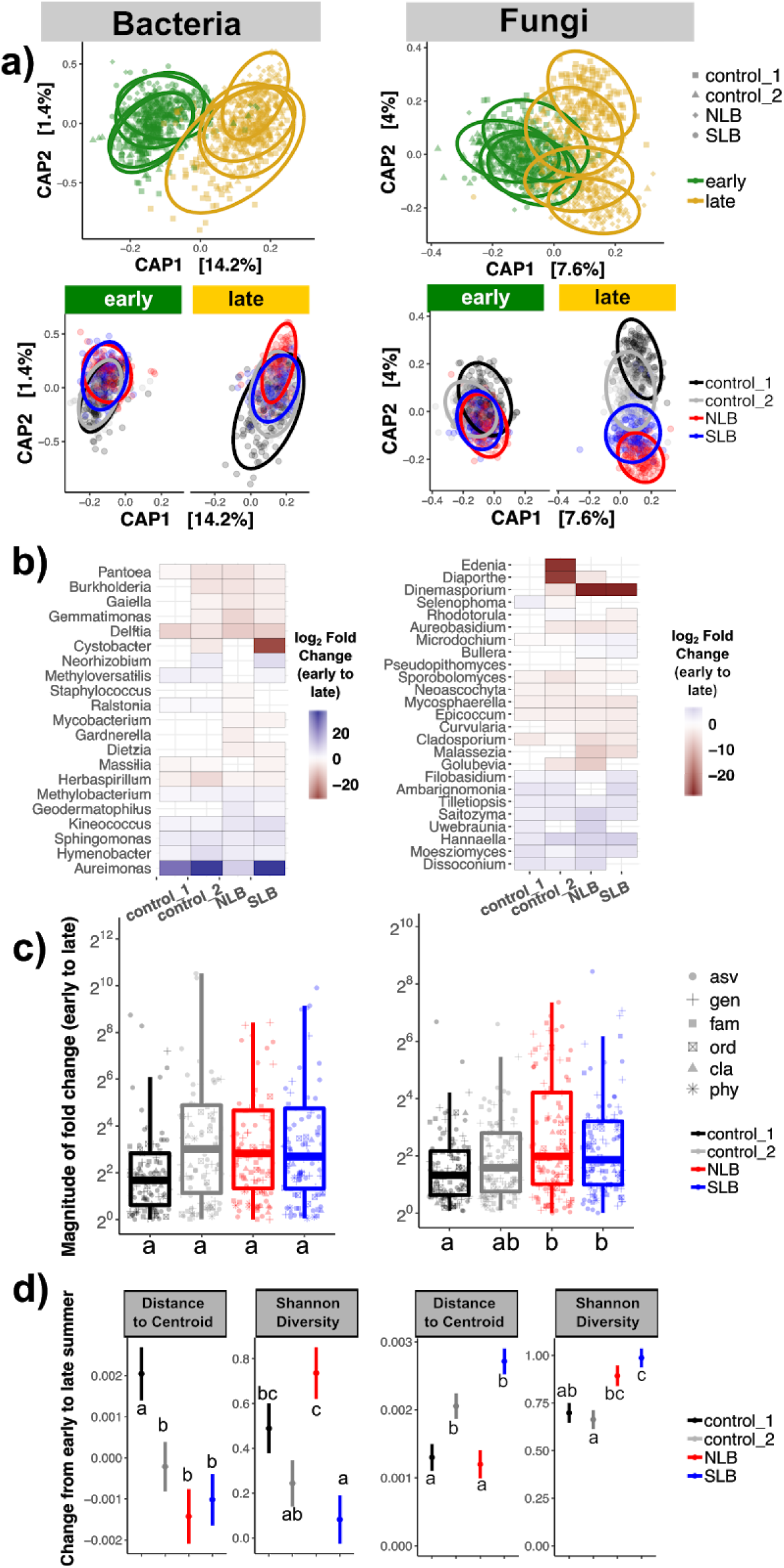
Maize leaf microbiomes (bacteria, left; and fungi, right) changed over the growing season regardless of disease pressure. All results are shown for four fields that are labeled according to the disease treatment they received (NLB/northern leaf blight, SLB/southern leaf blight, or none/control). **(a)** Partial distance-based redundancy analysis, constrained on the interaction between timepoint and field, shows that microbial communities in different fields became more distinct from each other over time. **(b)** Changes in relative abundance over time varied among fields for all taxa shown (likelihood ratio tests of negative binomial models with and without Timepoint*Field interaction term; FDR < 0.05). **(c)** The average magnitude of fold changes in relative abundance over time was similar between infected and non-infected fields. Fields with different letters were significantly different based on post-hoc Tukey tests, P < 0.05. Each point represents one taxon. The top, middle, and bottom lines of the boxes mark the 75th percentile, median, and 25th percentile, respectively; box whiskers extend 1.5 times the interquartile range above and below the box. The y-axis was truncated for clarity, obscuring several outliers. **(d)** Changes in alpha diversity (Shannon metric) and beta diversity (Distance to centroid) between timepoints differed among fields, but without regard to disease status (ANOVA, all P < 0.05; post-hoc Tukey tests, P < 0.05).

Because our disease treatments had to be applied to entire fields, replication was low and treatment was confounded with other factors such as the species of crops planted in adjacent fields, proximity to roads and trees, and the species of crops planted the previous year (Fig. 2; Table S2). As an additional test to circumvent this problem, we investigated whether temporal changes in microbiome composition and diversity were correlated with disease susceptibility within individual plants. We found no evidence that symptom severity altered microbiome succession in either NLB-infected or SLB-infected plants (Fig. S7). This result suggests that overall infection severity (measured at the whole-plant level) does not necessarily alter microbiome composition in the remaining green leaf tissue.

## Discussion

Breeding for MDR involves selecting alleles with the ability to alter the invasion success of several different pathogens. We demonstrated that different maize genotypes, identical except for the presence of QTL introgressed from two disease-resistant lines, assemble different leaf microbial communities both early in development and later in the growing season (Fig. 4; Fig. S6; Table 1; Table S6). This shift in community composition involved a wide variety of microbial taxa. Interestingly, some of these taxa (*e*.*g*., *Uwebraunia, Cryptococcus, Pseudopithomyces*) responded similarly to multiple independent introgressions (Fig. 4), suggesting that the underlying genes may involve partially redundant mechanisms. Many others, however, were consistently depleted in one field or timepoint but consistently enriched in a different environmental context (*e*.*g*., *Buchnera, Roseateles, Selenophoma, Moesziomyces*). Counterintuitively, in some environments seedlings of MDR genotypes were enriched in two fungal genera known to contain many plant pathogens (*Curvularia* and *Mycosphaerella*; Fig. 4a), although all plants were asymptomatic.

The inconsistency of these QTL effects among environments highlights one of the primary obstacles to understanding the relationship between host genotype and microbiome composition. Genotype-environment interactions for microbiome composition are strong and frequently observed (Agler et al. 2016; Peiffer et al. 2013; Wagner et al. 2016), contributing to the typically low heritability of these complex communities. Environmental variation has compounded effects on plant microbiomes because it not only directly influences the composition of the ambient pool of free-living organisms from which the host-associated community is derived, but also alters the expression of host genes and the emergent host phenotype (Lundberg et al. 2012; Wagner et al. 2016). This, in turn, determines the habitat available to potential symbionts. In general, these genotype-environment interactions greatly limit our ability to predict microbiome responses to changes in the host genotype, and therefore are a high-priority topic for future study (Busby et al. 2017). In the particular case of our study, they suggest that (1) disease resistance is not a reliable predictor of microbiome composition and (2) microbiome alteration is unlikely to be a mechanism through which MDR alleles confer improved disease resistance.

Nevertheless, our results are consistent with the hypothesis that some MDR loci may also affect non-pathogenic bacteria and fungi in certain environments. However, because these NILs carried introgressions covering up to 10% of the genome, we cannot rule out the possibility that linked genes--rather than the MDR alleles themselves--caused the observed shifts in microbiome composition. To determine whether the MDR alleles themselves caused the observed changes, follow-up experiments would need to compare the NILs with improved disease resistance to other NILs from the same population that did *not* show improved disease resistance. This, combined with data from a wider range of MDR lines, would greatly help to clarify the relationship between MDR *per se* and microbiome composition. Nevertheless, our results demonstrate that QTL introgression from disease-resistant lines can alter the relative abundances of diverse leaf symbionts (Fig. 4). Whether caused by linkage or true pleiotropy, these side-effects have the potential to either facilitate or interfere with the process of breeding for increased MDR (Fig. 1b).

We also hypothesized that in addition to directly affecting leaf microbiomes, MDR alleles would also indirectly influence them through cascading effects of improved disease resistance (Fig. 1a). For this reason, we expected to observe stronger host genotype effects after disease establishment. However, our data only partially supported this hypothesis, which relied on the assumption that disease establishment would profoundly disrupt the microbiome. This assumption was contradicted by our comparisons of microbiome composition in infected versus uninfected fields, and of severely versus mildly infected plants (Fig. 6; Fig. S7). We propose several possible explanations for the weaker-than-expected effect of pathogen invasion on the leaf microbiome. First, we deliberately sampled green tissue and avoided lesions (Fig. S2), which likely biased our dataset away from capturing the most strongly perturbed local communities. This choice was intentional because our primary interest was in direct effects of QTL introgression on non-pathogenic microbes; nevertheless, we expected to observe changes in microbiome composition as a result of the plant’s systemic response to infection (Gu *et al*., 2016; Hacquard *et al*., 2017). Second, the observed succession between timepoints likely reflected many different causal factors, including plant development and strong morphological differences between juvenile and adult leaves, a changing biotic context including insect communities and neighboring plants, and higher temperatures and humidity. The combined impact of these factors on the microbiome may have swamped out any signal of pathogen invasion. Finally, because our disease treatments could only be applied at the whole-field level, differences in microbial succession among fields also could have masked community responses to disease. A follow-up experiment that randomizes disease treatments while minimizing environmental variation would test this hypothesis.

Our finding that the introduced pathogens did not trigger strong cascading effects on the rest of the microbiome was surprising. One possible explanation is that other organisms were acting as keystone or “hub” taxa that interact with a large number of other microbes within the community (Agler *et al*. 2016; Herren & McMahon 2018). If such keystone taxa were insensitive to the presence of the pathogen, they may have had a stabilizing effect on the rest of the community. Keystone taxa also may have contributed to the highly variable effects of QTL introgression among environments. For example, it is possible that different taxa occupied “hub” positions in the microbial interaction networks within different environments, and that some of these hub organisms responded to the introgressed QTL while others did not. Improved statistical methods for analyzing microbial interaction networks, combined with manipulative experiments with synthetic microbial communities, would help to investigate this possibility (Vorholt *et al*. 2017; Röttjers & Faust 2018; Carr *et al*. 2019). Our results (e.g., Fig. 4) could reflect either direct effects of introgressed QTL or indirect effects via interacting microbes (Hassani *et al*. 2018).

Altogether, our results indicate that MDR can be improved in maize through introgression of QTL from disease-resistant lines, without major side-effects on microbiome structure or diversity. In our experiment such side effects were environment-specific and were limited to individual taxa (Fig. 4). The upshot for plant health--and ultimately, breeding outcomes--depends on whether individual symbionts increase or decrease in frequency during breeding, and whether they have a positive or negative effect on the host (Fig. 1b). The amplicon sequencing approach provides insufficient resolution to determine what effects the enriched or depleted taxa had on our experimental plants, if any. Re-inoculation experiments under controlled conditions would be necessary to determine whether these organisms affect disease resistance either positively or negatively. Another unresolved question that our data could not address is whether the introgressed QTL affected leaf microbiomes in ways other than changing relative abundance-- for example, by altering the total microbial load in leaves or by inducing changes in microbial gene expression and metabolic activity, which also could contribute to disease resistance (Chapelle *et al*., 2016). Understanding these complex links between the plant microbiota, pathogens, host phenotype, and environment will be crucial for developing microbiome-based solutions for sustainable disease control (Massart *et al*., 2015; Busby *et al*., 2017; Berg *et al*., 2017).

## Acknowledgements

We thank G. Marshall, J. Roberts, S. Sermons, and J. Torres for assistance with field and lab work, and we thank E. Barge, L. Burghardt, and D. Leopold for valuable comments and feedback on the manuscript. We thank Cathy Herring and the staff at NC Central Crops Research farm for managing the field trials. M.R.W. was supported by a NSF National Plant Genome Initiative Postdoctoral Research Fellowship in Biology (IOS-1612951). P.E.B. was supported by the NSF Science, Engineering, and Education for Sustainability Fellows program (CHE-1314095). This work was funded by a NCSU Plant Soil Microbial Community Consortium grant to P.B.K. and M.R.W.

## Author contributions

M.R.W., P.E.B., and P.B.K. designed the experiment. M.R.W. performed experiments and analyzed data. M.R.W. wrote the manuscript with contributions from P.E.B. and P.B.K.

---

**Table S1 |** Information on parentage of 11 maize genotypes used in this experiment.

**Table S2 |** Environmental and treatment information for the 5 fields used in this experiment.

**Table S3 |** The 20 most abundant genera in leaves of 3-week-old and 7-week-old maize plants.

**Table S4 |** Results of ANOVA of alpha diversity (ACE metric) for fungal and bacterial communities.

**Table S5 |** Results of ANOVA of beta diversity (distance to centroid) for fungal and bacterial communities.

**Figure S1 |** The introgressed QTL carried by the eight NILs in this study had little overlap.

**Figure S2 |** For diseased plants, we avoided lesions and targeted green tissue for microbiome analysis.

**Figure S3 |** The foliar fungal microbiome of three-week-old maize seedlings changed between years.

**Figure S4 |** Approximately one-third of all fungal ASVs in maize seedlings changed in relative abundance between 2016 and 2017.

**Figure S5 |** Effects of introgressed QTL on maize leaf alpha diversity (ACE and Shannon metrics).

**Figure S6 |** Effects of introgressed QTL on maize leaf beta diversity (distance to centroid).

**Figure S7 |** Within individual plants, temporal changes in community composition and alpha diversity did not correlate with disease resistance.

## Supporting Information

**Table S1.** Maize lines used in this experiment. The eight “DRIL” lines were near-isogenic lines descended from crosses between the disease-susceptible inbred line H100 and one of two disease-resistant inbred lines (Ki3 or NC304; Fig. 2a). On average, these DRIL lines retained ∼10% of their genome from the MDR donor line (Fig. S1).

| Line | MDR parent (donor) | MDS parent (recurrent) |
| --- | --- | --- |
| H100 | --- | H100 |
| Ki3 | Ki3 | --- |
| NC304 | NC304 | --- |
| DRIL32.063 | Ki3 | H100 |
| DRIL32.095 | Ki3 | H100 |
| DRIL32.134 | Ki3 | H100 |
| DRIL32.140 | Ki3 | H100 |
| DRIL62.030 | NC304 | H100 |
| DRIL62.032 | NC304 | H100 |
| DRIL62.054 | NC304 | H100 |
| DRIL62.127 | NC304 | H100 |

**Table S2.** Fields used for this experiment. All were located at Central Crops Research Station in Clayton, North Carolina, USA (Fig. 2). No pair of fields was separated by more than 2 km.

| Field | Soil type | Rotation (2014-2017) | Data collection | Disease Treatment (2017 only) |
| --- | --- | --- | --- | --- |
| A5B | Wagram loamy sand | Squash > Soybean > Corn > Corn | 2016-2017 | Uninoculated control |
| C10 | Dothan loamy sand | Soybean > Corn > Cotton > Corn | 2017 | SLB |
| D3 | Gilead sandy loam | Wheat/Fava bean > Cotton > Corn/Winter pea > Corn | 2017 | NLB |
| G4C | Norfolk loamy sand | Corn > Soybean > Corn > Tobacco | 2016 | NA |
| G5C | Wagram loamy sand | Corn > Tobacco > Cucumber > Corn | 2017 | Uninoculated control |

**Table S3.**
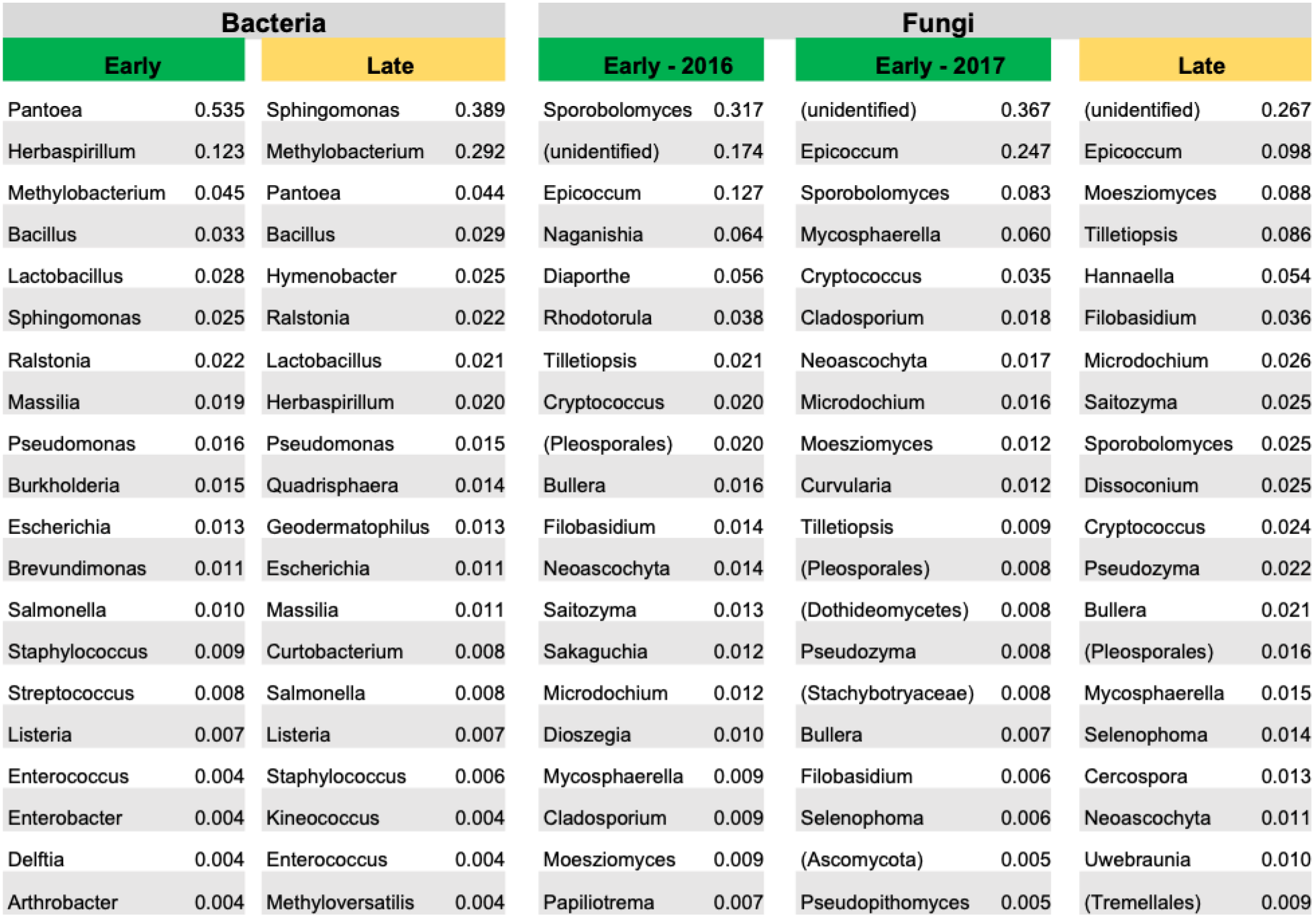
The 20 most abundant genera in leaves of 3-week-old (“early”) and 7-week-old (“late”) maize plants, with their relative abundances. Parentheses indicate groups that could not be identified at the genus level.

**Table S4.** Results of ANOVA of alpha diversity (ACE metric) for fungal and bacterial communities in the leaves of **(a)** maize seedlings three weeks after planting, and **(b)** adult maize seven weeks after planting. Linear mixed-effects models were fitted to log-transformed ACE values with predictors Genotype*Replicate while controlling for sequencing depth and batch effects; separate models were fit for the early and late timepoints. Least-squares mean estimates for each MDR genotype (relative to H100) are displayed in Fig. S5.

| (a) Early timepoint |  | Fungi |  | Bacteria |  |
| --- | --- | --- | --- | --- | --- |
|  |  | *F* test | *P* | *F* test | *P* |
| Genotype | | $F_{10,531} = 1.21$ | 0.28 | $F_{10,477} = 1.47$ | 0.15 |
| Rep | | $F_{5,38} = 25.91$ | $2.7e^{-11}$ | $F_{3,477} = 6.36$ | 0.00031 |
| Genotype x Rep | | $F_{48,527} = 1.68$ | 0.0038 | $F_{30,476} = 1.43$ | 0.070 |
| (b) Late timepoint |  | Fungi |  | Bacteria |  |
|  |  | *F* test | *P* | *F* test | *P* |
| Genotype | | $F_{10,542} = 1.68$ | 0.081 | $F_{10,495} = 2.63$ | 0.004 |
| Rep | | $F_{3,542} = 2.90$ | 0.034 | $F_{3,496} = 9.02$ | $8.0e^{-6}$ |
| Genotype x Rep | | $F_{30,542} = 1.33$ | 0.12 | $F_{30,495} = 2.21$ | 0.0003 |

**Table S5.** Results of ANOVA of beta diversity (distance from centroid) for fungal and bacterial communities in the leaves of **(a)** maize seedlings three weeks after planting, and **(b)** adult maize seven weeks after planting. Linear mixed-effects models were fitted to each individual’s distance to centroid, with predictors Genotype*Replicate while controlling for sequencing depth and batch effects; separate models were fit for the early and late timepoints. LS mean estimates for each MDR genotype (relative to H100) are displayed in Fig. S6.

| (a) Early timepoint | Fungi |  | Bacteria |  |
| --- | --- | --- | --- | --- |
|  | F test | P | F test | P |
| Genotype | $F_{10,855} = 1.42$ | 0.17 | $F_{10,536} = 1.37$ | 0.19 |
| Rep | $F_{5,60} = 23.08$ | $7.8e^{-13}$ | $F_{3,537} = 8.16$ | $2.5e^{-5}$ |
| Genotype x Rep | $F_{50,855} = 1.34$ | 0.063 | $F_{30,534} = 2.12$ | 0.0006 |
| (b) Late timepoint | Fungi |  | Bacteria |  |
|  | F test | P | F test | P |
| Genotype | $F_{10,555} = 3.19$ | 0.0005 | $F_{10,509} = 0.93$ | 0.50 |
| Rep | $F_{3,555} = 7.99$ | $3.2e^{-5}$ | $F_{3,509} = 33.93$ | $5.2e^{-20}$ |
| Genotype x Rep | $F_{30,555} = 2.33$ | 0.0001 | $F_{30,508} = 1.33$ | 0.11 |

**Table S6** | Negative binomial models identified dozens of microbial taxa that were differentially abundant in one or more MDR lines relative to the disease-susceptible parent line H100. Due to significant Genotype*Replicate interactions, estimates of log2-fold-changes differ among Replicates. Standard errors are provided for estimates of log2-fold-changes. Data from the early and late timepoints were analyzed separately. *P*-values were adjusted to correct for multiple comparisons using the false discovery rate (Benjamini & Hochberg, 1995). The relative abundance of each taxonomic group across the full dataset during the timepoint in question is also provided. This table is provided as a separate file in tab-delimited format.

**Figure S1.**
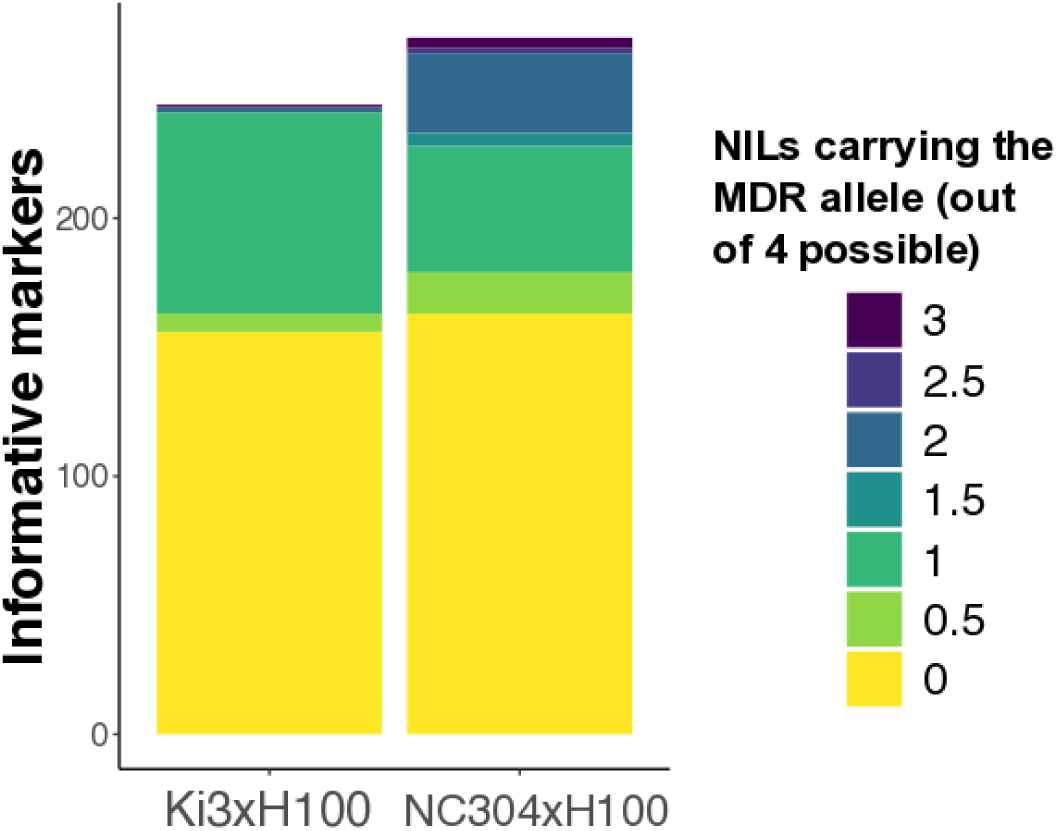
The introgressed MDR alleles carried by the eight NILs in this study had little overlap. The parent lines and NILs were genotyped using 245 or 270 informative markers (for the Ki3 x H100 cross and the NC304 x H100 cross, respectively). Most alleles from the MDR parent were present in only one NIL or not at all; for both crosses, all four NILs carried the H100 allele for >60% of markers. Observations of MDR alleles in the heterozygous state were scored as 0.5 rather than 1. Data from Lopez-Zuniga et al. (2019).

**Figure S2.**
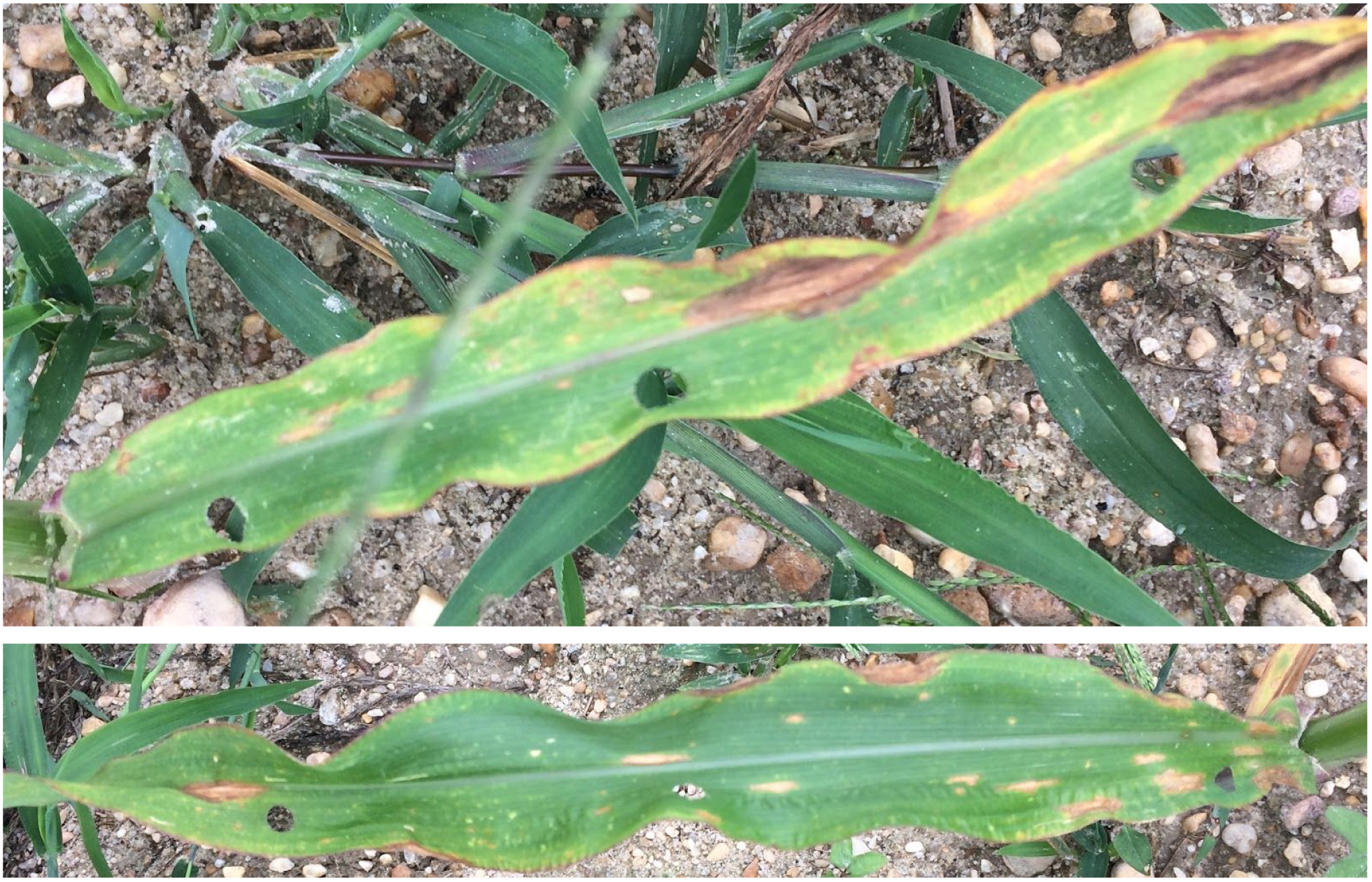
For diseased plants, we avoided lesions and targeted green tissue for microbiome analysis.

**Figure S3.**
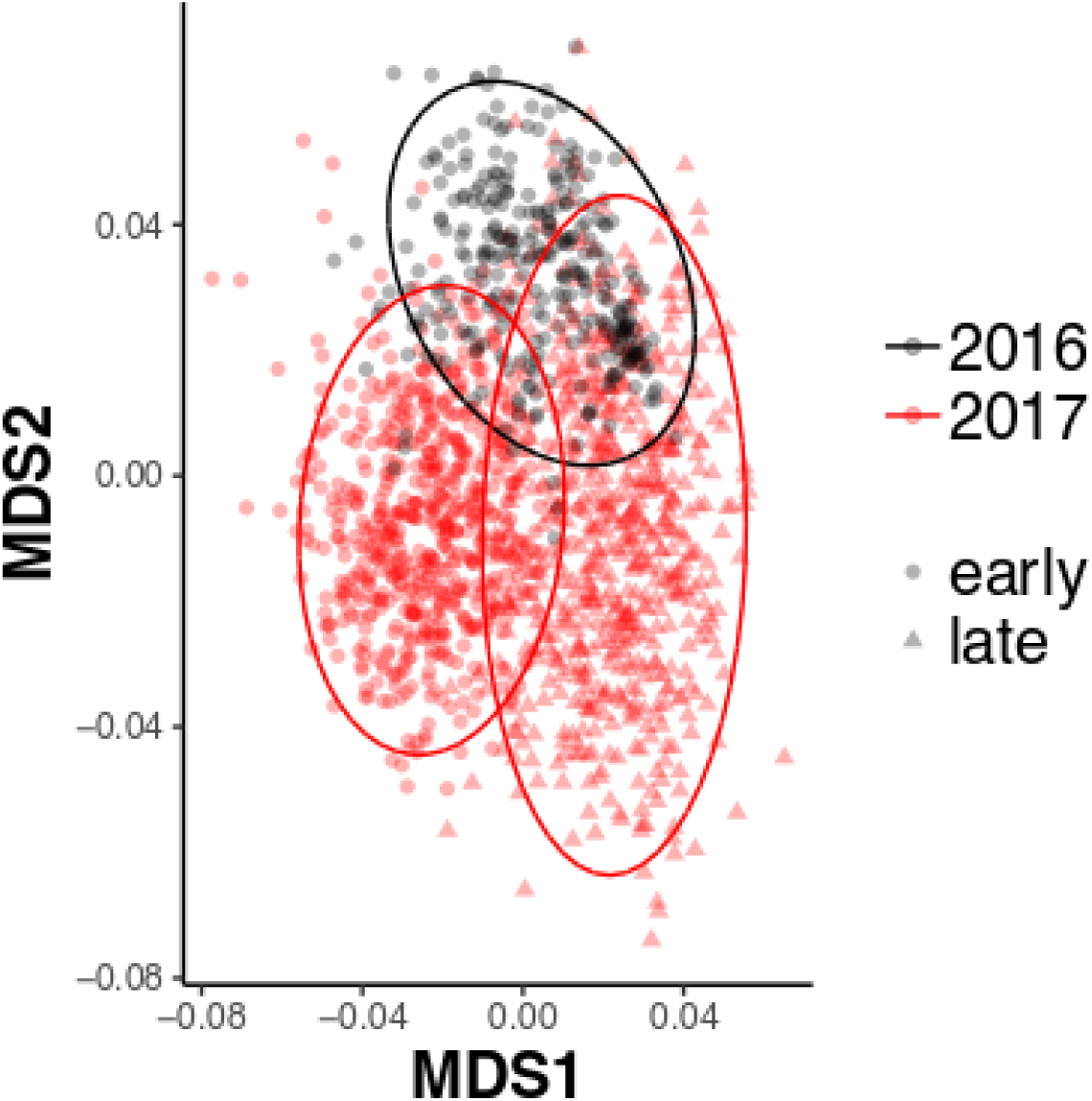
The foliar fungal microbiome of three-week-old maize seedlings changed between years. Non-metric multidimensional scaling of Bray-Curtis dissimilarities reveals that the early-timepoint samples from 2016 (black circles) cluster apart from the early-timepoint samples from 2017 (red circles); however, all early-timepoint samples (black and red circles) cluster apart from late-season samples (red triangles). Partial distance-based redundancy analysis was used to remove the effects of sequencing depth variation prior to ordination.

**Figure S4.**
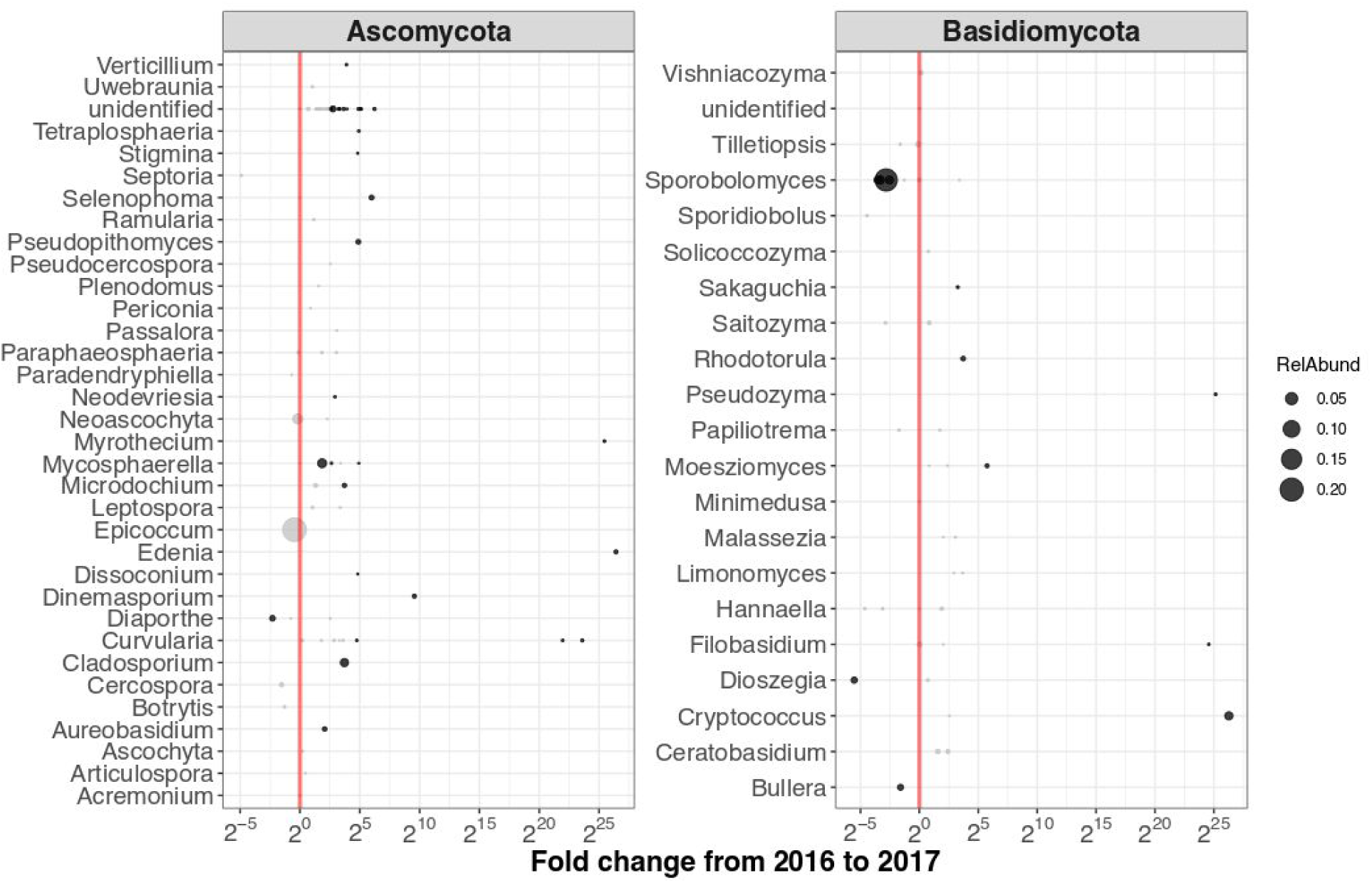
In seedlings growing in field “A5B”, the only field that was sampled in both years, many ASVs changed in relative abundance from 2016 to 2017. Grey points represent ASVs that did not change significantly between years; black points represent ASVs that were either more or less abundant in 2017 relative to 2016 (shown to the right or left of the dashed red line, respectively; FDR < 0.05). The area of each point is scaled by the relative abundance of the ASV that it represents.

**Figure S5.**
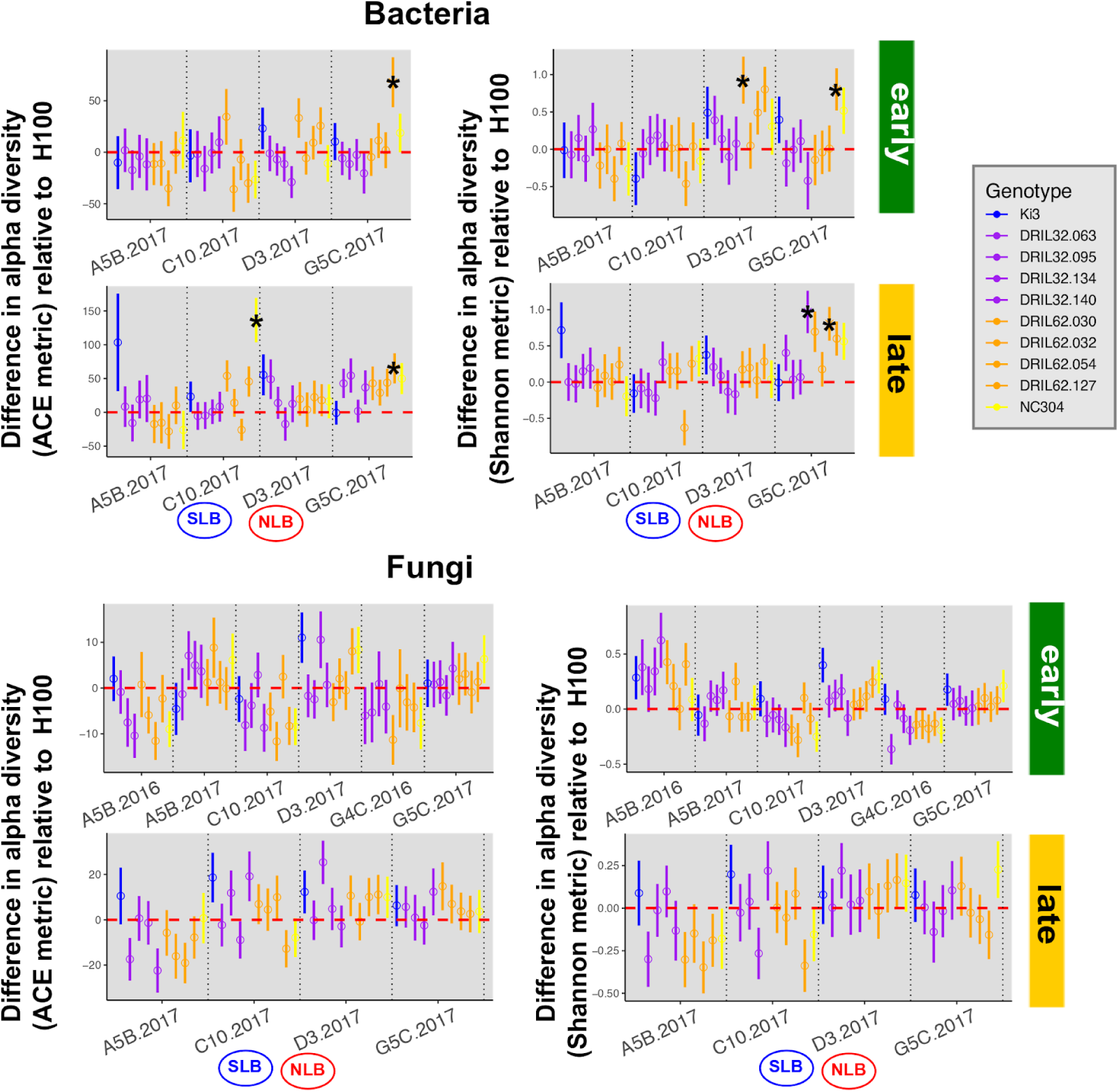
Introgression of MDR alleles shaped maize leaf alpha diversity (ACE and Shannon metrics). For each MDR genotype, its estimated deviation from the disease-susceptible line H100 is shown (based on LS means from linear mixed-effects models with predictors Genotype, Replicate, and Genotype*Replicate). Positive values for a given genotype indicate that within-sample diversity was higher than it was for the disease-susceptible line H100; negative values indicate that within-sample diversity was lower relative to H100. Error bars = +/- 1 s.e.m. Open circles mark deviations from H100 that were not significantly different from zero after *P*-value correction using Dunnett’s procedure; significant deviations from H100 are shown as asterisks. Corresponding ANOVA results are given in Table S4.

**Figure S6.**
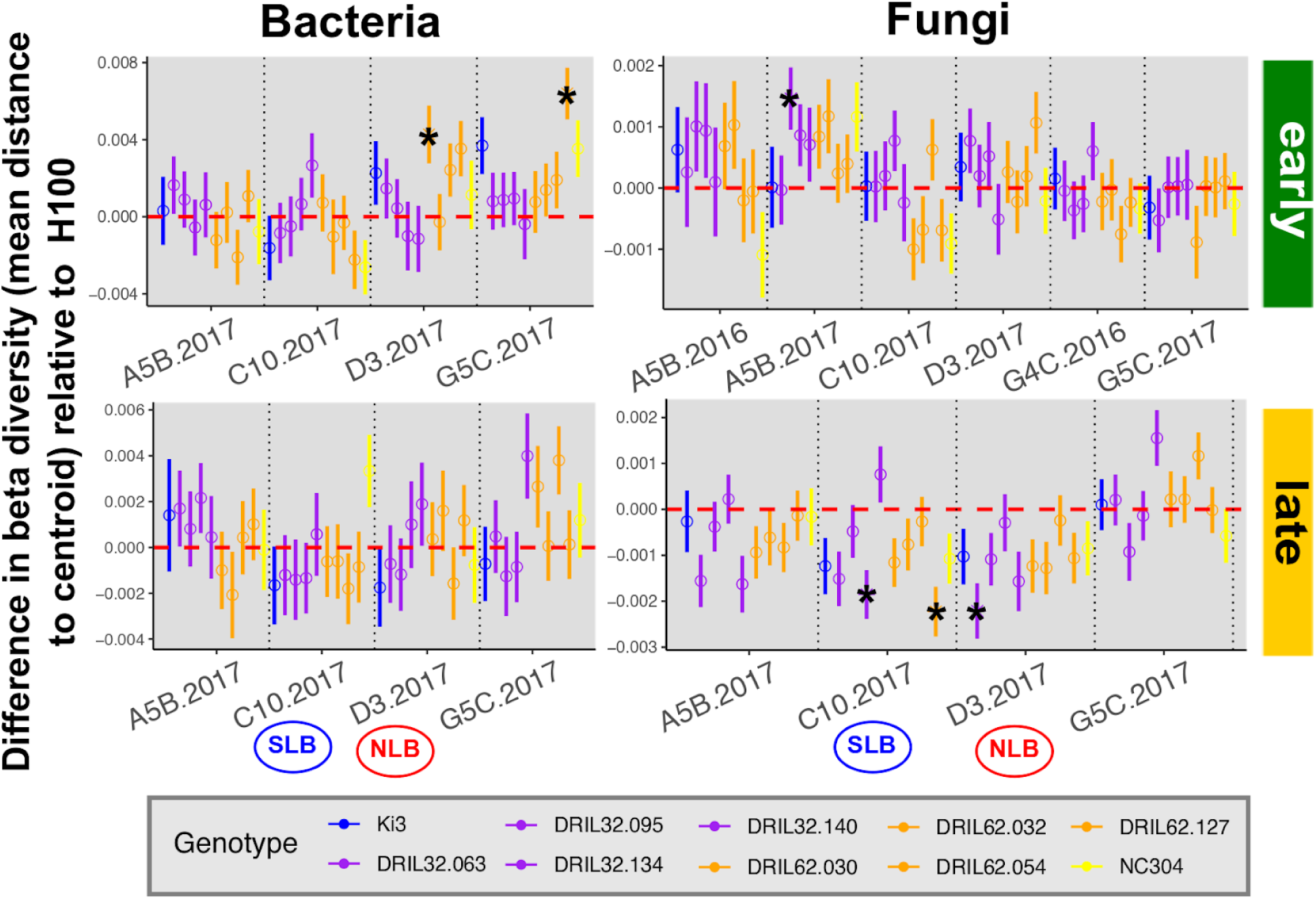
Introgression of MDR alleles altered maize leaf beta diversity (mean Distance to Centroid). For each MDR genotype, the average Distance to Centroid is shown in relation to the disease-susceptible line H100 is shown (based on LS means from linear mixed-effects models with predictors Genotype, Replicate, and Genotype*Replicate). Positive values indicate that inter-individual variation within a genotype was higher than it was within the disease-susceptible line H100; negative values indicate that inter-individual variation was lower than in H100. Error bars = +/- 1 s.e.m. Open circles mark deviations from H100 that were not significantly different from zero after *P*-value correction using Dunnett’s procedure; significant deviations from H100 are shown as asterisks. Corresponding ANOVA results are given in Table S4.

**Figure S7.**
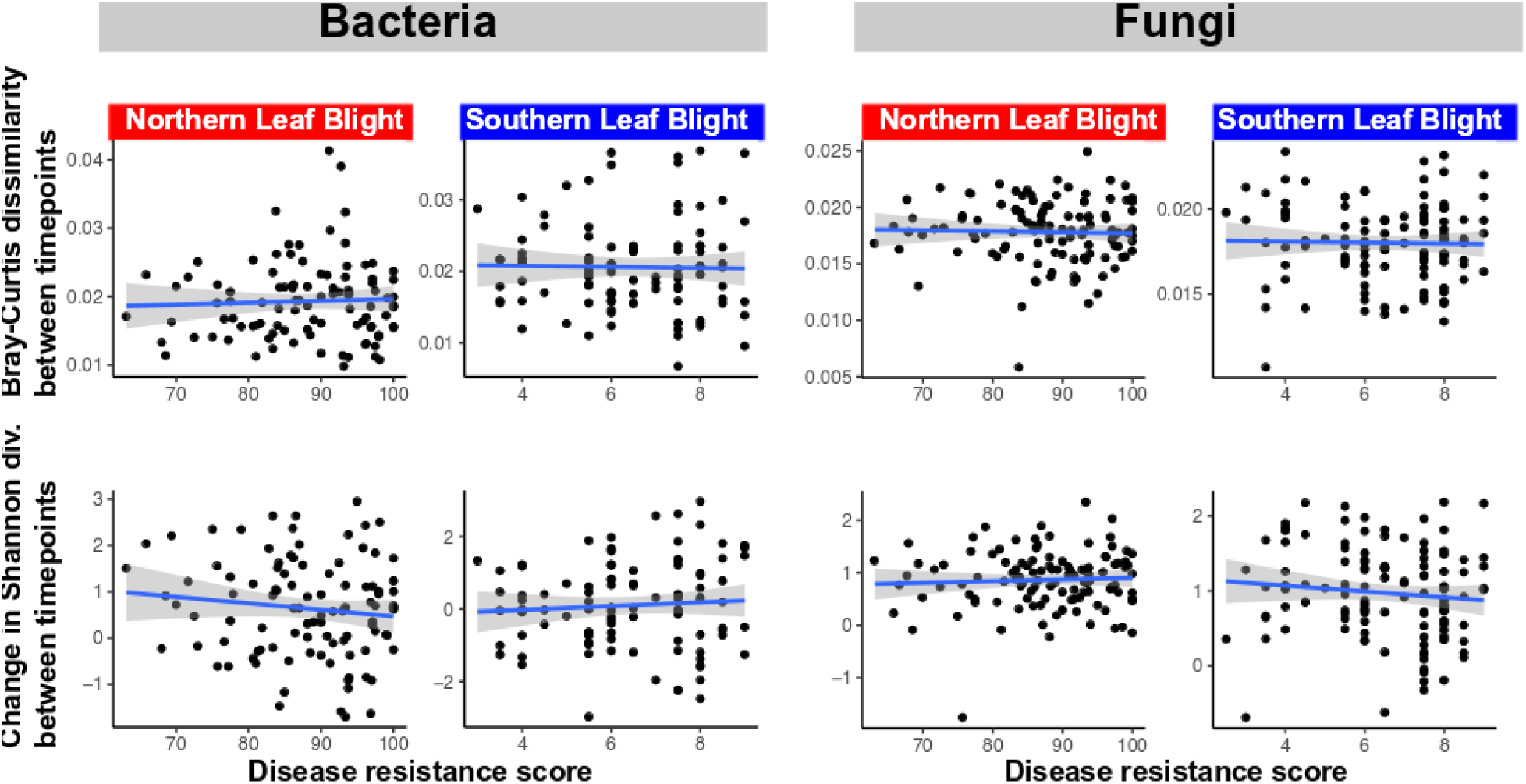
Within individual plants, temporal changes in community composition (top, quantified as Bray-Curtis dissimilarity between timepoints) and alpha diversity (bottom, quantified as the change in Shannon diversity between timepoints) did not correlate with disease resistance (linear regression, *P* > 0.05 for all tests). For NLB, *N* = 115; for SLB, *N* = 118.

